# Age and Sex Effects on Advanced White Matter Microstructure Measures in 15,628 Older Adults: A UK Biobank Study

**DOI:** 10.1101/2020.09.18.304345

**Authors:** Katherine E. Lawrence, Leila Nabulsi, Vigneshwaran Santhalingam, Zvart Abaryan, Julio E. Villalon-Reina, Talia M. Nir, Iyad Ba Gari, Alyssa H. Zhu, Elizabeth Haddad, Alexandra M. Muir, Emily Laltoo, Neda Jahanshad, Paul M. Thompson

**Affiliations:** Imaging Genetics Center, Mark and Mary Stevens Neuroimaging & Informatics Institute, University of Southern California, Marina del Rey, CA, USA

**Keywords:** aging, sex differences, white matter, diffusion-weighted MRI, microstructure

## Abstract

A comprehensive characterization of the brain’s white matter is critical for improving our understanding of healthy and diseased aging. Here we used diffusion-weighted magnetic resonance imaging (dMRI) to estimate age and sex effects on white matter microstructure in a cross-sectional sample of 15,628 adults aged 45-80 years old (47.6% male, 52.4% female). Microstructure was assessed using the following four models: a conventional single-shell model, diffusion tensor imaging (DTI); a more advanced single-shell model, the tensor distribution function (TDF); an advanced multi-shell model, neurite orientation dispersion and density imaging (NODDI); and another advanced multi-shell model, mean apparent propagator MRI (MAPMRI). Age was modeled using a data-driven statistical approach, and normative centile curves were created to provide sex-stratified white matter reference charts. Participant age and sex substantially impacted many aspects of white matter microstructure across the brain, with the advanced dMRI models TDF and NODDI detecting such effects the most sensitively. These findings and the normative reference curves provide an important foundation for the study of healthy and diseased brain aging.

## Introduction

White matter alterations have been linked to age-related cognitive decline and implicated in neurodegenerative diseases such as Alzheimer’s disease (Bennett & Madden, 2014; Pievani, Filippini, van den Heuvel, Cappa, & Frisoni, 2014). A range of age-associated neurodegenerative diseases also exhibit sex differences in their prevalence and presentation and, similar to age, sex has also been associated with white matter differences (Cox et al., 2016; Jahanshad & Thompson, 2017; Ritchie et al., 2018; Salminen, Tubi, Bright, & Thompson, 2020; Toschi, Gisbert, Passamonti, Canals, & De Santis, 2020). Understanding the brain’s white matter may substantially improve our understanding of aging and sex differences therein, including ultimately the genetic and environmental factors that may influence healthy or diseased aging. Diffusion-weighted magnetic resonance imaging (dMRI) allows for the characterization of white matter microstructure by assessing the diffusion of water molecules in brain tissue (Stejskal & Tanner, 1965). The conventional modeling approach applied to dMRI data, known as diffusion tensor imaging (DTI), fits a single-tensor to single-shell dMRI data and typically reflects hindered diffusion (Basser, Mattiello, & Lebihan, 1994; Jones, 2008). A more advanced single-shell model is the tensor distribution function (TDF), which addresses well-established limitations of DTI by using a continuous mixture of tensors to capture multiple underlying fiber populations (Leow et al., 2009; Nir et al., 2017; Zhan et al., 2009). Compared to single-shell models, multi-shell dMRI models may allow for a more nuanced depiction of the underlying microstructural environment by using multi-shell dMRI data, which allows both hindered and restricted diffusion to be captured. Multi-shell diffusion models include, among others, the biophysical model neurite orientation dispersion and density imaging (NODDI) and the signal-based model mean apparent propagator MRI (MAPMRI). NODDI is a multi-compartment model that separately models restricted, hindered, and free water diffusion, which are thought to correspond to intra-cellular, extra-cellular, and isotropic water components, respectively (Zhang, Schneider, Wheeler-Kingshott, & Alexander, 2012); NODDI may thus provide microstructure metrics more closely linked to specific aspects of the cellular environment than single-shell models (Zhang et al., 2012), although some recent work suggests the assumptions underlying NODDI’s specificity may not always be met (Jelescu & Budde, 2017; Jelescu, Palombo, Bagnato, & Schilling, 2020). MAPMRI is a diffusion propagator-based multi-shell model that estimates the diffusion patterns of water molecules without *a priori* assumptions about the underlying tissue, which may allow for the detection of more subtle microstructure alterations (Fick, Wassermann, Caruyer, & Deriche, 2016; Le et al., 2020; Ning et al., 2015; Ozarslan et al., 2013).

Previous dMRI studies examining age and sex effects have reported age-related white matter decline and significant sex differences in white matter microstructure (Beck et al., 2020; Cox et al., 2016; Damoiseaux, 2017; Jahanshad & Thompson, 2017; Ritchie et al., 2018; L.E. Salminen et al., 2020; Toschi et al., 2020; Tseng et al., 2020; Zavaliangos-Petropulu et al., 2019). In one of the largest studies to date that investigated age and sex associations with white matter microstructure adults, Cox et al. (2016) examined two DTI metrics (fractional anisotropy, FA; mean diffusivity, MD) and three NODDI metrics (orientation dispersion, OD; intra-cellular volume fraction, ICVF; isotropic volume fraction, ISOVF) in 3,513 middle-aged and older subjects from the UK Biobank for a range of white matter tracts across the brain; age was modeled using a linear or quadratic fit, based on the best fit for each tract and metric. These analyses indicated widespread effects of age and sex on most tracts for the examined DTI and NODDI metrics; interactions between age and sex exhibited small effect sizes and attained statistical significance for a limited subset of the tracts and metrics examined (Cox et al., 2016). A larger follow-up investigation in 5,216 UK Biobank participants assessed sex differences in one DTI metric (FA) and one NODDI metric (OD), among other non-dMRI measures, and similarly supported the existence of significant white matter differences between men and women for a range of tracts (Ritchie et al., 2018). In another study among 7,167 UK Biobank participants, Tseng et al. (2020) examined age effects on white matter using a linear age model and four microstructure metrics analogous to FA or calculated using DTI (generalized fractional anisotropy, GFA; mean diffusivity, MD; axial diffusivity, AD; radial diffusivity, RD). Their results further supported the association between age and white matter decline, in addition to indicating a sex difference in the overall number of white matter tracts that exhibited age-related declines in anisotropy (Tseng et al., 2020). Such prior dMRI work robustly demonstrates that multiple aspects of white matter microstructure are significantly associated with participant age and sex (Beck et al., 2020; Cox et al., 2016; Damoiseaux, 2017; Jahanshad & Thompson, 2017; Ritchie et al., 2018; Salminen et al., 2020; Toschi et al., 2020; Tseng et al., 2020; Zavaliangos-Petropulu et al., 2019). However, it remains an open question how age, sex, and their interaction may be related to additional measures of white matter microstructure obtained from other advanced dMRI models such as TDF and MAPMRI, as well as how age effects on microstructure may manifest in middle to late adulthood when using more complex, data-driven statistical approaches for modeling age.

Our understanding of aging would also benefit from establishing typical ranges of white matter properties among middle-aged and older adults. Such normative reference data would allow future investigations to detect individuals with quantifiably abnormal white matter microstructure for their age and sex, such as individuals who are below the 5^th^ percentile or above the 95^th^ percentile for a given microstructure metric (Marquand et al., 2019; Wolfers et al., 2018; Zabihi et al., 2019). As many age-related neurodegenerative diseases are associated with altered white matter, such normative white matter reference curves may allow future aging work to identify individuals with the most severe neural pathology (Pievani et al., 2014). This would provide a novel avenue for future pre-clinical aging studies by allowing them to characterize those factors associated with the most diseased neural phenotypes. Prior large-scale neuroimaging has computed normative reference values for a range of structural morphometry measures, such as hippocampal volume, as well as several microstructure metrics (Dima et al., 2020; Frangou et al., 2020; Nobis et al., 2019; Pomponio et al., 2020; Tseng et al., 2020). Normative reference curves for additional white matter microstructural properties would provide complementary information and, together with preexisting neural reference values, allow for a more complete characterization of healthy and diseased aging in future studies.

Here we expanded on prior work by using multiple dMRI models to thoroughly characterize age and sex effects on white matter microstructure in a large-scale, population-based sample of middle-aged and older adults. Specifically, we examined age and sex associations with DTI, TDF, NODDI, and MAPMRI microstructure metrics in 15,628 cross-sectional UK Biobank participants. Age was modeled non-linearly using a data-driven statistical approach, and normative centile curves were calculated for all dMRI measures to provide sex-stratified references for white matter. We found that age and participant sex was significantly related to many white matter properties across the brain, and advanced dMRI models detected age and sex effects the most sensitively. The computed reference curves provide a novel avenue for future studies focused on the characterization of white matter in middle to late adulthood.

## Methods

### Study Design, MRI Acquisition and Processing

We analyzed cross-sectional dMRI data from a total of 15,628 community-based UK Biobank subjects aged 45-80 years (47.6% male) (Miller et al., 2016). All dMRI data included in the current study was collected on a single scanner. Sample details and dMRI processing are presented in the Supplementary Material (Supplementary Methods). Briefly, white matter metrics were derived using four dMRI reconstruction models: DTI, TDF, NODDI, and MAPMRI. Each reconstruction method is further described in the Supplementary Material (Supplementary methods). Metrics derived from DTI included fractional anisotropy (FA^DTI^), mean diffusivity (MD), axial diffusivity (AD), and radial diffusivity (RD). An advanced measure of fractional anisotropy was calculated using TDF (FA^TDF^). Measures derived from NODDI included orientation dispersion (OD), intra-cellular volume fraction (ICVF), and isotropic volume fraction (ISOVF). The following white matter indices were calculated from MAPMRI: return-to-origin probability (RTOP), return-to-axis probability (RTAP), and return-to-plane probability (RTPP). Diffusion-weighted MRI metrics were projected to a standard white matter skeleton using publicly available ENIGMA protocols based on FSL’s tract-based statistics (TBSS) and described further in the Supplementary Material (Supplementary Methods) (http://enigma.ini.usc.edu/protocols/dti-protocols; Jahanshad et al., 2013; Smith et al., 2006). Consistent with prior literature, we focused on mean microstructure values for the whole white matter skeleton (full WM) and the corpus callosum (CC) (e.g., Beck et al., 2020; Jahanshad & Thompson, 2017; Pines et al., 2020). For completeness, supplemental analyses examined mean values in additional white matter regions across the brain (Supplemental Methods).

### Statistical Analyses

Our planned *a priori* analyses investigated the effects of age, sex, and their interaction on the full WM and CC by using fractional polynomials to flexibly model age in a non-linear manner (Dima et al., 2020; Frangou et al., 2020; Royston & Altman, 1994). The fractional polynomial approach is detailed further in the Supplementary Material (Supplementary Methods), as are the nuisance covariates we included based on prior literature (Salminen et al., 2019). Effect sizes were calculated as the variance explained separately by age, sex, and their interaction. For instance, the effect size for age was computed as the difference in variance (change in R^2^) between two models: one which included age in addition to sex and nuisance covariates, and one which only included sex and nuisance covariates. Sex-stratified centile reference curves were created for each white matter region and dMRI metric using quantile regression (Supplementary Methods). For completeness, supplemental analyses examined age and sex effects on additional white matter regions across the brain, in addition to using a distinct statistical approach to model age; details on these supplemental analyses are presented in the Supplementary Material (Supplementary Methods).

## Results

Age was robustly associated with the full WM and CC for all dMRI metrics (Fig. 1A-B, Fig. 2, Fig. 3). Such age effects in our cross-sectional sample were observed in the full WM and CC as *lower* anisotropy (FA^DTI^, FA^TDF^), neurite density (ICVF), and restriction (RTOP, RTAP, RTPP), as well as *higher* diffusivity (MD, AD, RD) and free water (ISOVF) with increasing age. White matter dispersion (OD) was higher in the full WM and lower in the CC with increasing age. Effect sizes for age in the full WM and CC demonstrated that the advanced single-shell model, TDF, was most sensitive to age, followed by DTI and NODDI (Fig. 1A–B). Supplemental analyses likewise supported that TDF was the most sensitive dMRI model, followed by DTI (Supplementary Results). In sum, age has widespread effects on white matter microstructure that are detected most sensitively by TDF.

**Figure 1.**
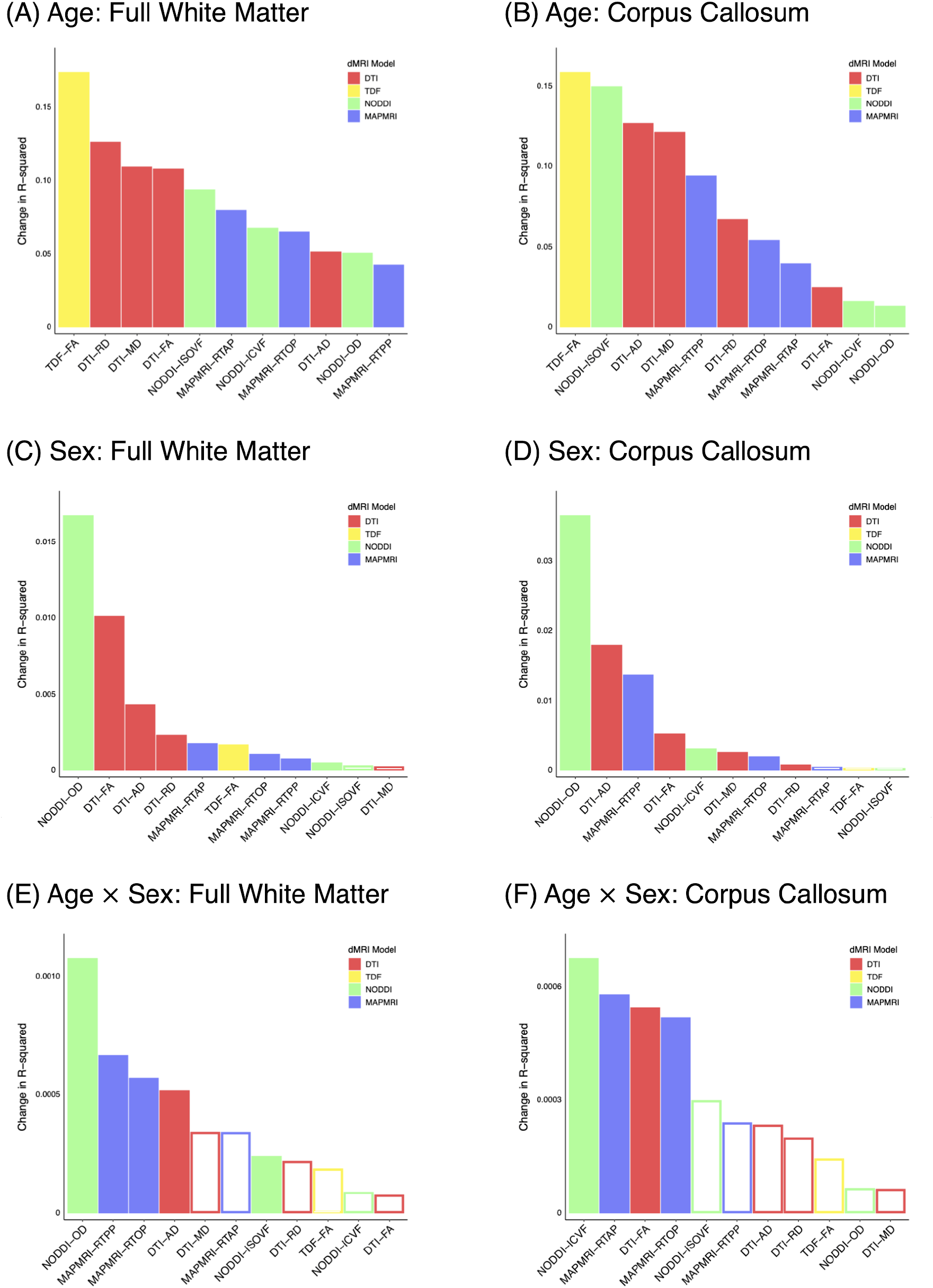
Effect of age (A-B), participant sex (C-D), and their interaction (E-F) on full white matter and corpus callosum microstructure. Age was modeled as a continuous variable using fractional polynomials. Filled bars indicate a significant association, whereas hollow bars indicate the association did not attain statistical significance.

**Figure 2.**
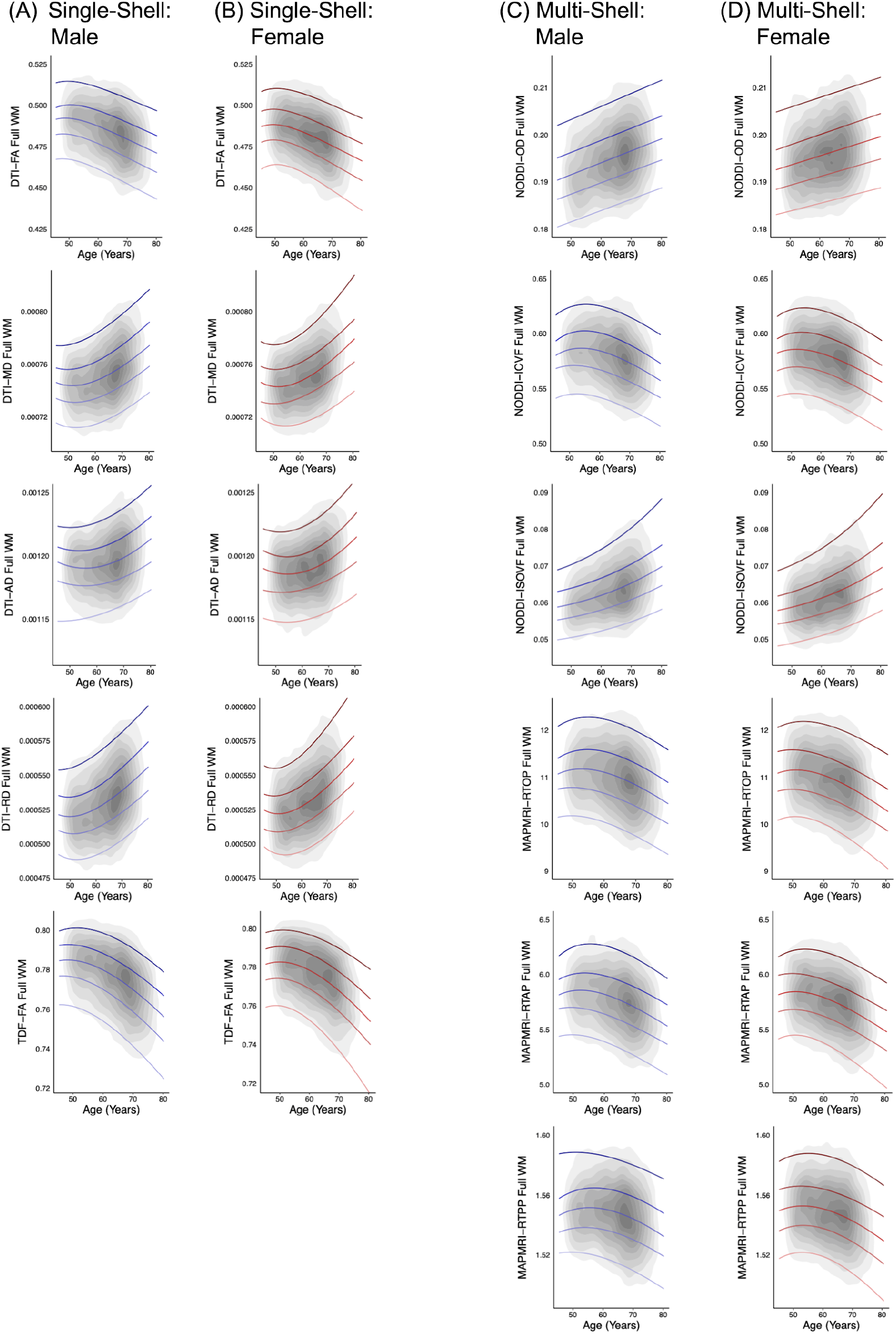
Normative centile reference curves calculated for the full white matter for single-shell dMRI metrics in (A) males and (B) females, and multi-shell dMRI metrics in (C) males and (D) females. Solid colored lines, ordered from lightest to darkest, indicate the following centiles: 5^th^, 25^th^, 50^th^, 75^th^, 95^th^; blue lines indicate male participants, and red lines indicate female participants. Gray overlay reflects kernel density (darker=greater degree of data point overlap). Full WM = full white matter skeleton.

**Figure 3.**
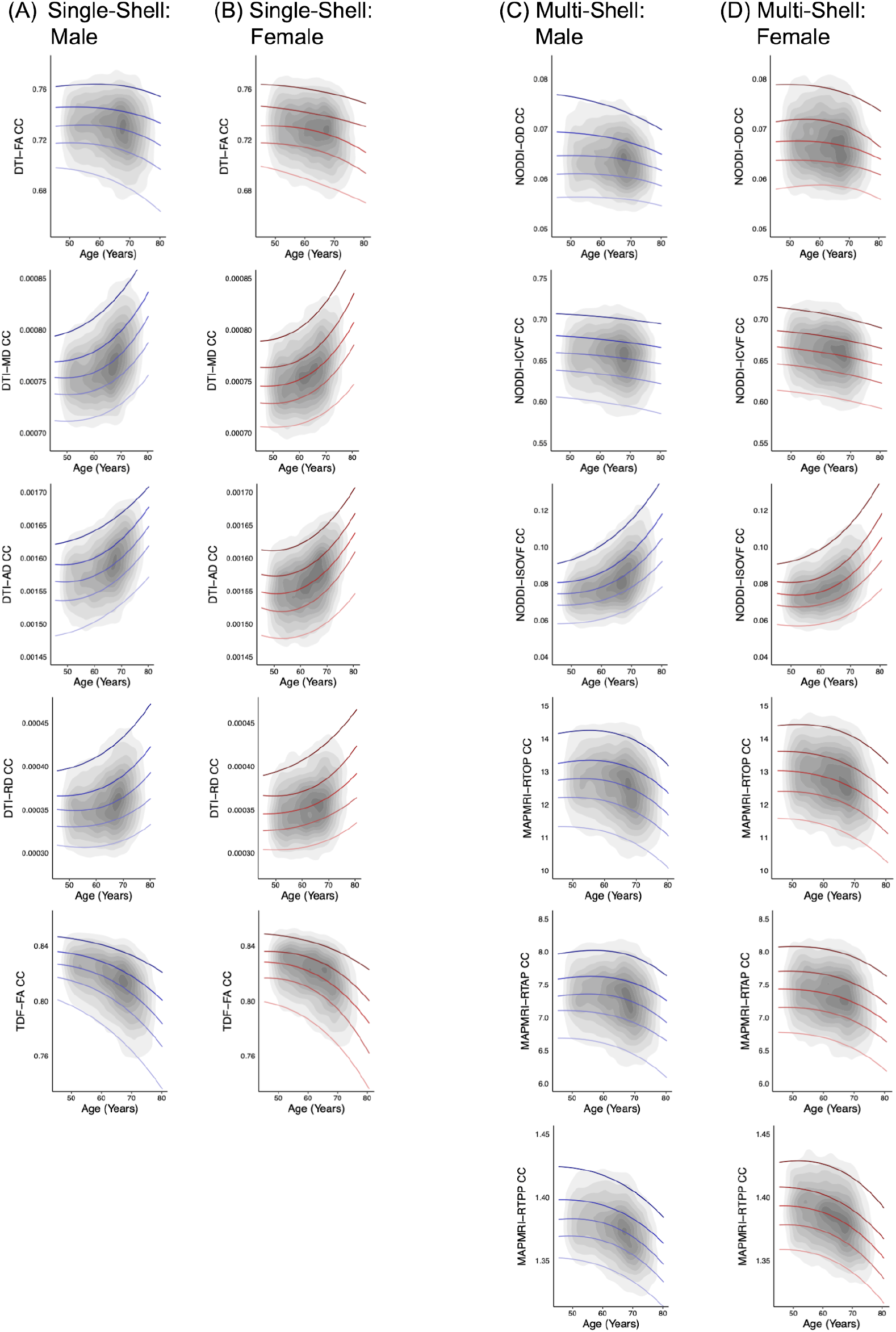
Normative centile reference curves calculated for the corpus callosum for single-shell dMRI metrics in (A) males and (B) females, and multi-shell dMRI metrics in (C) males and (D) females. Solid colored lines, ordered from lightest to darkest, indicate the following centiles: 5^th^, 25^th^, 50^th^, 75^th^, 95^th^; blue lines indicate male participants, and red lines indicate female participants. Gray overlay reflects kernel density (darker=greater degree of data point overlap). CC = corpus callosum.

Participant sex was significantly associated with full WM and CC microstructure for nearly all assessed dMRI indices (Fig. 1C-D, Fig. 2, Fig. 3). Patterns of sex differences were typically region- and metric-specific. In both the full WM and the CC, women exhibited greater RD, OD, and RTPP than men; men displayed greater FA^DTI^ and AD compared with women. For full WM, male participants also displayed higher FA^TDF^, ICVF, RTOP, and RTAP. When contrasting CC microstructure between men and women, women had greater ICVF and RTOP. Men exhibited higher MD in the CC than women. Effect sizes for full WM and CC revealed that sex differences were detected most sensitively by the advanced multi-shell model, NODDI, followed by DTI (Fig. 1C-D). Supplemental analyses similarly demonstrated that NODDI and DTI were most sensitive to sex differences (Supplementary Results). Together, these results indicate that participant sex is robustly associated with white matter across the brain, and such sex differences are captured most sensitively by NODDI.

Age effects on the full WM and CC significantly depended on participant sex for multiple dMRI metrics (Fig. 1E-F, Fig. 2, Fig. 3). These age by sex interactions in our cross-sectional sample were driven by complex combinations of steeper age-related decline among women than men and vice versa. The most common pattern was generally steeper decline in female than male participants, as observed in both the full WM (AD, ISOVF, RTPP) and the CC (ICVF, RTOP). A mixture of faster, slower, and similar age-associated decline in female compared to male subjects was also observed to a somewhat lesser extent, in both full WM (RTOP) and the CC (FA^DTI^, RTAP). In full WM only, men displayed steeper age-related decline than women in fiber dispersion (OD). Effect sizes for the age by sex interaction in the full WM and CC indicated that the multishell model, NODDI, was most sensitive to such sex differences in age effects, followed by MAPMRI and DTI (Fig. 1E-F). Supplemental analyses similarly indicated that NODDI and DTI exhibited the greatest effect sizes (Supplementary Results). As a whole, the effect of age significantly depends on participant sex for many white matter regions and measures, and NODDI detects such sex differences in age effects the most sensitively.

To provide normative models of white matter microstructure for the full WM and CC, sex-stratified centile reference curves were calculated for each dMRI metric (Fig. 2, Fig. 3). For completeness, normative reference charts for additional white matter ROIs across the brain are included in the Supplementary Material (Supplementary Results).

## Discussion

Here we thoroughly characterized age and sex effects on white matter microstructure by using an array of dMRI models coupled with advanced statistical methods. We also created normative reference curves for multiple aspects of white matter micro-architecture, which may allow for the future identification of individuals with the greatest neural pathology. As a whole, age and participant sex was robustly related to white matter across the brain, with the advanced dMRI models TDF and NODDI capturing such differences the most sensitively.

Age was associated with white matter alterations in nearly every region and microstructural property investigated. Specifically, older age in our cross-sectional sample was related to mostly *lower* anisotropy, neurite density, and restriction, as well as mostly *higher* diffusivity and free water; age-dependent changes in fiber dispersion were more regionally specific. These results are in line with postmortem histological findings that aging is associated with the degradation and deformation of axons and myelination (Bennett & Madden, 2014). Our age results are also consistent with prior dMRI work that examined one or two dMRI models among the DTI, TDF, NODDI, and MAPMRI approaches included here (Beck et al., 2020; Bennett & Madden, 2014; Cox et al., 2016; Damoiseaux, 2017; Tseng et al., 2020; Zavaliangos-Petropulu et al., 2019). Notably, white matter microstructure in aging has previously been linked to age-related cognitive decline and neurodegenerative diseases, underscoring the importance of understanding age effects on white matter among older adults (Bennett & Madden, 2014; Pievani et al., 2014).

Participant sex was also significantly related to many white matter properties across the brain. As a whole, women exhibited greater white matter dispersion than men, on average. Men displayed greater anisotropy, on average, compared to women. Sex differences in other white matter microstructure characteristics – including measures reflecting diffusivity, neurite density, free water, and restriction – depended to a greater extent on the exact region and metric examined. These findings are largely consistent with previous analyses which used DTI or NODDI to assess white matter sex difference (Cox et al., 2016; Jahanshad & Thompson, 2017; Ritchie et al., 2018; Salminen et al., 2020; Toschi et al., 2020); such prior analyses likewise found that males exhibit higher anisotropy and lower fiber dispersion than females, with other microstructure metrics displaying greater variability in their pattern of sex differences. In the current study, we also observed significant sex differences in age effects for a number of regions and microstructure metrics in our cross-sectional sample. The most common pattern for this interaction between age and sex was steeper age-related decline among women than men. Previous dMRI work has likewise suggested sex differences in age effects (Cox et al., 2016; Kodiweera, Alexander, Harezlak, McAllister, & Wu, 2016; Toschi et al., 2020; Tseng et al., 2020). The largest of these studies investigated 7,167 UK Biobank participants and found that, overall, a greater number of tracts exhibited age-related declines in anisotropy among women than men (Tseng et al., 2020). Another study in 3,513 subjects from the UK Biobank demonstrated that age effects significantly differed between men and women for a small number of tracts and microstructure metrics, where the directionality of such differences depended on the specific tract and metric (Cox et al., 2016). Our findings expand on this prior work indicating sex differences in age effects among middleaged to older adults. Importantly, a range of neurodegenerative conditions exhibit sex differences in their prevalence and presentation, emphasizing the importance of understanding differences in neural aging between men and women (Salminen et al., 2020).

Advanced dMRI models detected age and sex effects most sensitively in the current study. We used four separate dMRI models to characterize white matter microstructure – DTI, TDF, NODDI, and MAPMRI – and found that the advanced models TDF and NODDI exhibited the greatest sensitivity to white matter differences. Notably, between the two single-shell models DTI and TDF, TDF may be considered to model the underlying neurobiology more directly than DTI; among the two multi-shell models, NODDI may similarly be considered to model the underlying biology more directly than MAPMRI. More specifically, among the single-shell dMRI approaches, TDF models multiple underlying fibers per voxel, whereas DTI cannot differentiate multiple fiber populations (Jones, 2008; Leow et al., 2009; Nir et al., 2017; Zhan et al., 2009). Similarly, among the multi-shell dMRI models included in the current study, NODDI directly models multiple aspects of the cellular environment whereas MAPMRI estimates diffusion patterns without specifically modeling the underlying biology (Fick et al., 2016; Ozarslan et al., 2013; Zhang et al., 2012). This suggests that dMRI approaches which model the underlying neurobiology may capture microstructural differences more sensitively in community-based samples of middle-aged and older adults such as the UK Biobank sample examined here. As TDF was the most sensitive to age and NODDI to participant sex, our results furthermore indicate that the relative utility of each dMRI model depends not only on the fidelity of the model to the underlying neurobiology, but also on the specific neurobiology underlying the scientific question of interest. Future work should assess whether the relative utility of different dMRI models may depend on the dMRI analysis method used (e.g., TBSS vs. tractography-based approaches). As a whole, these findings indicate that future research examining age and sex may benefit from including advanced dMRI measures.

The age and sex findings presented here may serve as a reference for future analyses investigating the genetic and environmental factors that contribute to healthy or diseased aging. The normative centile charts provided here may also allow for the detection of individuals with abnormal white matter, contributing to future studies focused on the characterization of diseased aging. Future work should expand on the current study by examining longitudinal samples, replicating our findings in diverse healthy and diseased datasets collected on different scanners, and including additional dMRI analysis methods beyond the TBSS approach used here, such as tractography-based analyses.

## Conclusions

In summary, we characterized white matter microstructure during middle to late adulthood in 15,628 individuals using multiple dMRI models. Age and participant sex exhibited widespread associations with white matter across the brain, and advanced dMRI models demonstrated the greatest sensitivity to such effects. These findings provide an important foundation for the study of healthy and diseased aging.

## Acknowledgments

This research was conducted using the UK Biobank resource under Application 11559.

## Availability of data and material

Diffusion-weighted MRI data are available through the UK Biobank application procedure (https://www.ukbiobank.ac.uk/register-apply/).

## Code availability

Code for the ENIGMA dMRI protocol is publicly available here: http://enigma.ini.usc.edu/protocols/dti-protocols.

