## Supplementary Material for "Age and Sex Effects on Advanced White Matter Microstructure Measures in 15,628 Older Adults: A UK Biobank Study"

#### Supplementary Methods

*Participants*

The UK Biobank is a publicly available dataset of community-based middle-aged and older adults residing in the United Kingdom, described in greater detail elsewhere (Miller et al., 2016). In the current study, we analyzed available cross-sectional data from UK Biobank participants who had usable dMRI scans collected on a single scanner. Neuroimaging data collection for the UK Biobank is on-going, and all data included here was collected in Cheadle, Manchester prior to the COVID-19 pandemic. Subjects were only included in the current sample if their chromosomal sex was identical to their sex as recorded elsewhere in the UK Biobank (i.e., participant sex as reported by the subject or as recorded in the subject's medical records). Additionally, participants were required to have complete nuisance covariate data (see *Statistical Analyses*). Our final sample included data from a total of 15,628 UK Biobank subjects aged 45-80 years, with an approximately equal number of males and females (47.6% male, 52.4% female). Additional demographic information is presented in Table S1.

*MRI Acquisition and Processing*

Diffusion-weighted MRI scans were acquired and preprocessed as described previously (Alfaro-Almagro et al., 2018; Miller et al., 2016). Briefly, all dMRI images included in the current study were collected on a single Siemens Skyra 3T scanner with a 32-channel headcoil using a standard ("monopolar") Stejskal-Tanner pulse sequence and consisted of two diffusion-weighted shells ($b$=1000 and 2000 s/mm$^2$) with 50 distinct diffusion-encoding directions each, for a total of

100 directions (voxel dimensions = 2mm$^3$ isotropic, field of view = 104x104x72, acquisition time = 7 minutes). Preprocessing consisted of correction for eddy currents, head motion, and outlier slices using FSL's Eddy tool, followed by gradient distortion correction (Alfaro-Almagro et al., 2018; Andersson & Sotiropoulos, 2015; Andersson & Sotiropoulos, 2016). In the current study, white matter metrics were derived using four dMRI reconstruction models: DTI, TDF, NODDI, and MAPMRI. Each reconstruction method is further described below (see *Reconstruction Models*). For single-shell dMRI data ($b$=1000 s/mm$^2$, 50 diffusion-encoding directions), the conventional model, DTI, was fit by the UK Biobank using FSL's DTIFIT (Alfaro-Almagro et al., 2018). The advanced model, TDF, was fit in-house using custom scripts (Nir et al., 2017). Metrics derived from DTI included fractional anisotropy ($FA^{DTI}$), mean diffusivity (MD), axial diffusivity (AD), and radial diffusivity (RD). TDF was used to derive an advanced measure of fractional anisotropy ($FA^{TDF}$). For multi-shell dMRI data ($b$=1000 and 2000 s/mm$^2$, 100 diffusion-encoding directions total), the advanced models NODDI and Laplacian-regularized MAPMRI were fit by the UK Biobank and in-house using the AMICO tool and DIPY, respectively (Daducci et al., 2015; Fick, Wassermann, Caruyer, & Deriche, 2016; Garyfallidis et al., 2014). The following white matter indices were calculated using NODDI: orientation dispersion (OD), intra-cellular volume fraction (ICVF), and isotropic volume fraction (ISOVF). Measures derived from MAPMRI included return-to-origin probability (RTOP), return-to-axis probability (RTAP), and return-to-plane probability (RTPP).

Diffusion-weighted MRI metrics were projected in-house to a standard white matter skeleton using publicly available ENIGMA protocols (http://enigma.ini.usc.edu/protocols/dti-protocols; Jahanshad et al., 2013). Briefly, each subject's whole-brain $FA^{DTI}$ map was nonlinearly registered to the ENIGMA $FA^{DTI}$ template using ANTs's symmetric image normalization (SyN) method (Avants, Epstein, Grossman, & Gee, 2008), and the resulting transformation was applied to the maps for each metric. White matter metrics were projected onto the ENIGMA template skeleton based on the $FA^{DTI}$ metric using FSL's tract-based spatial statistics (TBSS; Smith et al.,

2006). Major white matter regions of interest (ROIs) were labeled in-house using the Johns Hopkins University white matter atlas (Mori et al., 2008). For our planned *a priori* analyses, mean values for each dMRI metric were extracted from the whole white matter skeleton (full WM) and from the largest white matter structure in the brain, the corpus callosum (CC), consistent with prior literature (e.g., Beck et al., 2020; Jahanshad & Thompson, 2017; Pines et al., 2020). For supplementary analyses, mean values for each metric were extracted from an additional 14 white matter ROIs (Table S2).

*Statistical Analyses*

Primary analyses investigated the effects of age, sex, and their interaction on the full WM and CC. We used fractional polynomials (*mfp* in R; R Core Team, 2016) to flexibly model age in a non-linear manner without specifying *a priori* the relationship between age and the dependent variable (Dima et al., 2020; Frangou et al., 2020; Royston & Altman, 1994). Briefly, the best-fitting age model was found for the full WM and CC for each dMRI metric separately by assessing one- and two-term curvilinear age models with the following possible powers (Dima et al., 2020; Frangou et al., 2020; Royston & Altman, 1994): -2, -1, -0.5, 0, 0.5, 1, 2 and 3, where $x^0$ corresponds to *ln(x)*. In line with prior work (Salminen et al., 2019), all analyses included the following nuisance covariates: educational attainment (operationalized as “college” or “no college”), socioeconomic status (quantified using the Townsend Deprivation Index; Townsend, Phillimore, & Beattie, 1988), waist-hip ratio, and population structure (measured using the first 4 principal components obtained from the UK Biobank’s genetic ancestry analysis; Bycroft et al., 2017). Effect sizes were calculated as the variance explained separately by age, sex, and their interaction. For instance, the effect size for age was computed as the difference in variance (change in $R^2$) between two models: one which included age in addition to sex and nuisance covariates, and one which only included sex and nuisance covariates.

For completeness, supplemental analyses characterized age and sex effects on an additional 14 white matter ROIs across the brain (Table S2) by using the statistical approach described above; a false discovery rate (FDR) of 5% was applied to correct for multiple comparisons across the 14 supplemental ROIs assessed (Benjamini & Hochberg, 1995). To confirm all results were robust to statistical approach, we followed previously published fractional polynomial recommendations to repeat analyses after discretizing the continuous variable of interest (Royston & Sauerbrei, 2004). For these supplemental analyses, we binarized age by splitting participants into two groups: $\geq$ 60 years old and < 60 years old; this binarization threshold was selected based on previous large-scale neuroimaging studies examining aging (Dima et al., 2020; Frangou et al., 2020).

Sex-stratified centile reference curves were created for each white matter region and dMRI metric. Specifically, these normative charts modeled the fractional polynomial age terms determined above by using quantile regression (*quantreg* in R) to calculate the following percentiles: 5, 25, 50, 75, and 95.

*Reconstruction Models*

**Diffusion Tensor Imaging (DTI)**. DTI models a single fiber orientation by fitting a single diffusion tensor, or ellipsoid, at each voxel of the brain in single-shell dMRI data (Basser, Mattiello, & Lebihan, 1994). DTI assumes a 3D Gaussian diffusion process, which is fit using six independent tensor parameters (3 eigenvalues and 3 eigenvectors) that together describe the shape and orientation of the tensor. Here we used DTI to create maps of tensor-derived fractional anisotropy ($FA^{DTI}$), which represents the degree of diffusion anisotropy within a fiber bundle (Jones, 2008).

$$FA = \sqrt{\frac{3}{2}} \frac{\sqrt{(\lambda_1 - <\lambda>)^2 + (\lambda_2 - <\lambda>)^2 + (\lambda_3 - <\lambda>)^2}}{\sqrt{\lambda_1^2 + \lambda_2^2 + \lambda_3^2}}$$

The eigenvalues are represented as $\lambda_1$, $\lambda_2$, $\lambda_3$, and the average diffusivity in all directions, or mean diffusivity (MD), is represented as $< \lambda >$. Axial diffusivity (AD) is defined as $\lambda_1$, the primary (largest) eigenvalue. Radial diffusivity (RD) is calculated as the average of $\lambda_2$ and $\lambda_3$, the second and third eigenvalues. In the current study, we assessed the metrics $FA^{DTI}$, MD, AD, and RD.

**Tensor Distribution Function (TDF)**. TDF models single-shell dMRI data by using a probabilistic mixture of tensors, which allows for the reconstruction of multiple underlying fiber populations, together with a distribution of weights (Leow et al., 2009; Nir et al., 2017). For each voxel, the TDF is calculated as the probability distribution function *P*(D(θ, λ))* of all feasible Gaussian tensors *D(θ, λ).* The tensor orientation distribution function (TOD) is then computed as the marginal density function of the TDF.

$$TOD(\theta) = \int_{\lambda} P(D(\theta, \lambda))\, d\lambda$$

Eigenvalues are calculated for each spherical angle θ by computing the expected value of each eigenvalue λ along θ. From this, the advanced fractional anisotropy measure $FA^{TDF}$ is calculated.

$$FA^{TDF} = \int TOD(\theta) * FA(\theta)\, d\theta$$

$$FA(\theta) = \frac{\sqrt{(\lambda'_1(\theta) - \lambda'_2(\theta))^2 + (\lambda'_1(\theta) - \lambda'_3(\theta))^2 + (\lambda'_2(\theta) - \lambda'_3(\theta))^2}}{2[\lambda'_1(\theta)^2 + \lambda'_2(\theta)^2 + \lambda'_3(\theta)^2]}$$

$$\lambda'_i(\theta) = \frac{\int P(D(\theta, \lambda))\lambda_i\, d\lambda}{\int P(D(\theta, \lambda))\, d\lambda}$$

In the current study, we included $FA^{TDF}$ in the examined dMRI metrics.

**Neurite Orientation Dispersion and Density Imaging (NODDI)**. NODDI is a multi-compartment model applied to multi-shell dMRI data that models the complete dMRI signal $E(\mathbf{q})$ as consisting of intra-cellular, extra-cellular, and isotropic water components; each of these components uniquely affects diffusion in the environment and results in a separate dMRI signal (Zhang, Schneider, Wheeler-Kingshott, & Alexander, 2012).

$$E(\mathbf{q}) = v_{iso}E_{iso}(\mathbf{q}) + (1 - v_{iso})[v_{ic}E_{ic}(\mathbf{q}) + v_{ec}E_{ec}(\mathbf{q})]$$

The dMRI signal of the intra-cellular, extra-cellular, and isotropic compartments are represented here as $E_{ic}$, $E_{ec}$, $E_{iso}$, respectively. The corresponding volume fractions are $v_{ic}$, $v_{ec}$, $v_{iso}$, where $v_{ec} = 1 - v_{ic}$ . $E_{ic}$ is modeled as a set of dispersed cylinders of zero radius, where the dispersion is defined by a Watson distribution. $E_{ec}$ represents a dispersed mixture of Gaussian anisotropic diffusion, and $E_{iso}$ represents isotropic Gaussian diffusion (Daducci et al., 2015; Zhang et al., 2012). In the current study, we investigated orientation dispersion (OD), intra-cellular volume fraction (ICVF), and isotropic volume fraction (ISOVF) (Daducci et al., 2015; Zhang et al., 2012).

**Mean Apparent Propagator MRI (MAPMRI)**. MAPMRI is a diffusion propagator-based model applied to multi-shell dMRI data that estimates water diffusion without requiring *a priori* assumptions regarding the underlying tissue (Fick et al., 2016; Ozarslan et al., 2013). Specifically, MAPMRI models the acquired dMRI signal $E(\mathbf{q})$ as a set of continuous orthogonal basis functions which together represent the space $E(\mathbf{q}; \mathbf{c})$. In this formulation, the diffusion signal is represented as the basis coefficients c and the q-space wave vector $\mathbf{q} = |\mathbf{q}|\mathbf{g}$ , where the gradient direction $\mathbf{g}$ is related to the b-value as $|\mathbf{q}| = \sqrt{b/(\Delta - \delta/3)}/2\pi$ . We describe the estimation of the basis coefficients **c** here, with the formulation of the MAPMRI basis functions detailed elsewhere (Fick et al., 2016). Briefly, the fitting of **c** is regularized such that $\hat{E}(\mathbf{q};\mathbf{c})$ uses Laplacian regularization to smoothly interpolate between the measured q-space points, and λ represents the regularization weight (Fick et al., 2016).

$$\mathrm{argmin}_{\mathrm{c}} \int_{\mathbb{R}^3} \left[E(\mathbf{q}) - \hat{E}(\mathbf{q};\mathbf{c})\right]^2 d\mathbf{q} + \lambda \int_{\mathbb{R}^3} \left[\nabla^2 \hat{E}(\mathbf{q};\mathbf{c})\right]^2 d\mathbf{q}$$

After **c** is determined, the MAPMRI basis represents both the dMRI signal and the diffusion propagator. In the current study, we considered the q-space indices return-to-origin probability (RTOP), return-to-axis probability (RTAP), and return-to-plane probability (RTPP) (Fick et al., 2016; Ozarslan et al., 2013).

#### Supplementary Results

When investigating age effects on dMRI indices, supplemental analyses examining additional white matter ROIs across the brain and using a distinct statistical approach to model age demonstrated that TDF was the most sensitive dMRI model, followed by DTI (Fig. S1A, Fig. S2-S15, Fig. S16A-B, Fig. S17A). These results are consistent with our primary analyses and further suggest that age has widespread effects on white matter microstructure which are detected most sensitively by TDF.

With regards to the impact of sex on microstructure, follow-up analyses investigating supplemental white matter ROIs and using different methods to model age similarly found that NODDI and DTI were most sensitive to sex differences (Fig. S1B, Fig. S2-S15,Fig. S16C-D, Fig. S17B). In line with our primary results, these findings indicate that participant sex is robustly associated with white matter across the brain, and that such sex differences are captured most sensitively by NODDI.

When examining the interaction between participant age and sex, supplementary analyses characterizing further WM ROIs and testing the robustness of results to the selected age model demonstrated that NODDI and DTI exhibited the greatest effect sizes (Fig. S1C, Fig. S2-S15, Fig. S16E-F, Fig. S17C). Similar to our main analyses, these results indicate that the impact of age significantly depends on participant sex for many white matter regions and measures, and that NODDI detects such interactions the most sensitively.

For completeness, sex-stratified centile reference curves were calculated for each dMRI metric for additional white matter ROIs across the brain to provide normative models of white matter aging (Fig. S2-S15).

### Supplementary Tables and Figures

**Table S1.** Mean and Standard Deviation of Sample Descriptives

| | Male | Female |
|---|---|---|
| Sample Size | 7442 | 8186 |
| Age (Years) | 63.6 ± 7.6 | 62.2 ± 7.3 |
| Waist/Hip Ratio | 0.92 ± 0.06 | 0.81 ± 0.06 |
| Socioeconomic Status (Townsend Index) | -2.1 ± 2.64 | -2.0 ± 2.6 |
| Education (College / No College) | 5388 / 2054 | 5154 / 3032 |

**Table S2.** White Matter Regions of Interest Analyzed from the Johns Hopkins University (JHU) Atlas

| Abbreviation | Tract Name |
|---|---|
| CGC | Cingulum (cingulate) |
| CGH | Cingulum (hippocampal) |
| CR | *Corona radiata* |
| CST | Corticospinal tract |
| EC | External capsule |
| FX | Fornix (body) |
| FXST | Fornix *(crus)* / *stria terminalis* |
| IC | Internal capsule |
| PTR | Posterior thalamic radiation |
| SFO | Superior fronto-occpital fasciculus |
| SLF | Superior longitudinal fasiculus |
| SS | *Sagittal stratum* |
| TAP | Tapetum |
| UNC | Uncinate fasciculus |

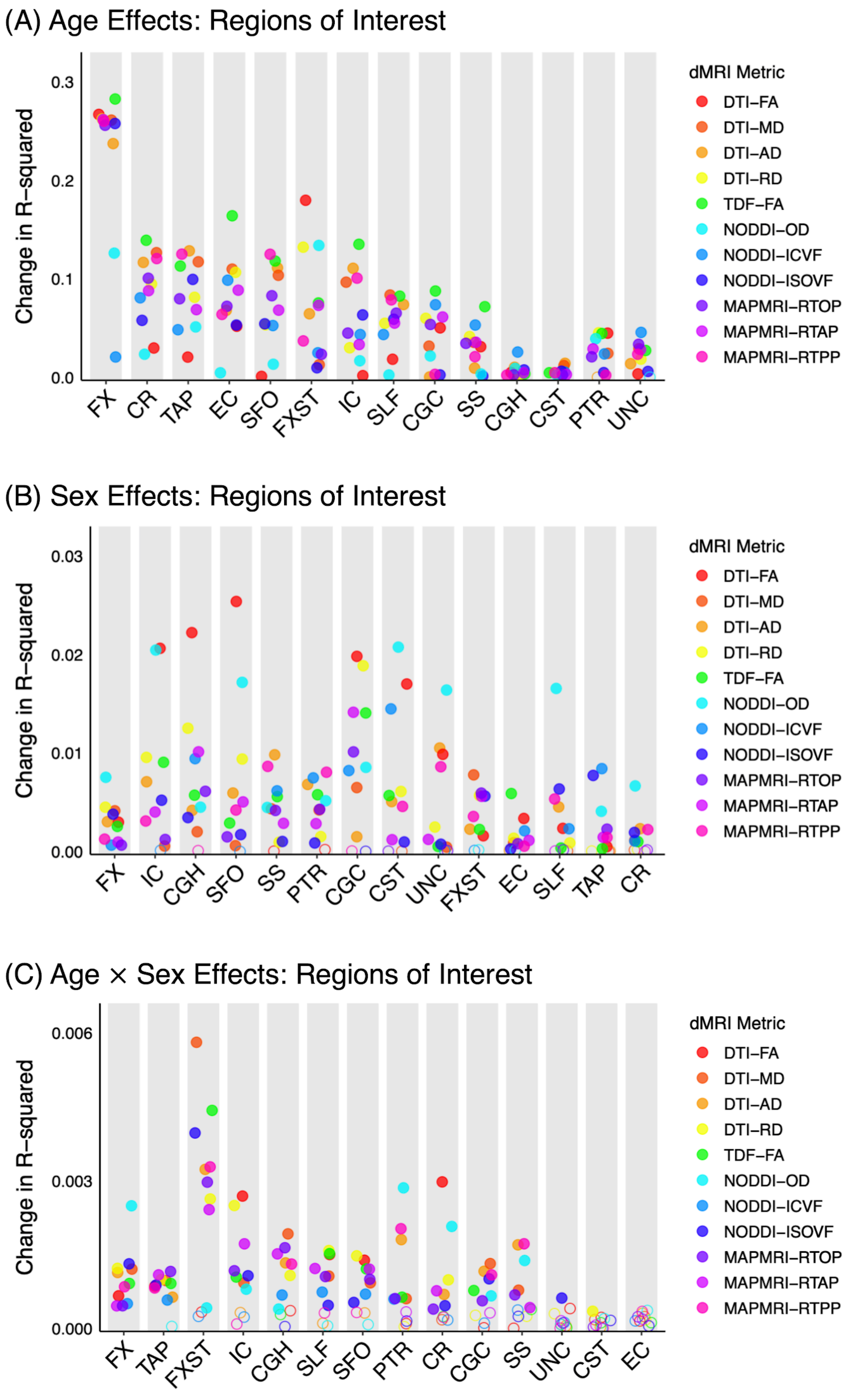


**Figure S1.** Effect of age (A), participant sex (B), and their interaction (C) on white matter microstructure when modeling age with fractional polynomials. Filled circles indicate the association was significant after FDR-correction for the number of regions. Regions are ordered by the number of significant metrics, followed by the mean effect size. CGC = cingulum (cingulate), CGH = cingulum (hippocampal), CR = *corona radiata*, CST = corticospinal tract, EC = external capsule, FX = fornix (body), FXST = fornix (*crus*) / *stria terminalis*, IC = internal capsule, PTR = posterior thalamic radiation, SFO = superior fronto-occpital fasciculus, SLF = superior longitudinal fasiculus, SS = *sagittal stratum*, TAP = tapetum, UNC = uncinate fasciculus.

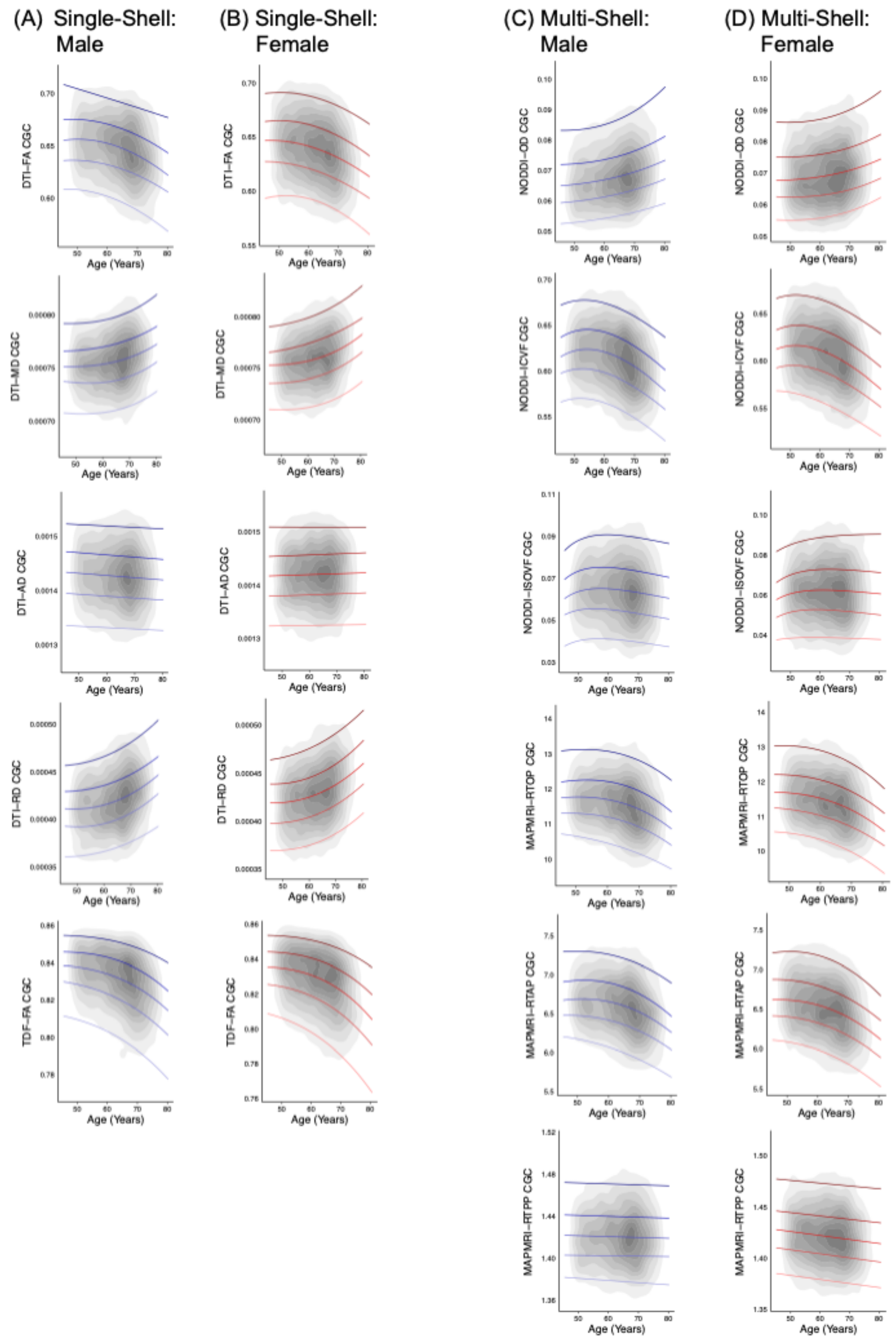


**Figure S2.** Normative centile reference curves calculated for the cingulum (cingulate) for single-shell dMRI metrics in (A) males and (B) females, and multi-shell dMRI metrics in (C) males and (D) females. Solid colored lines, ordered from lightest to darkest, indicate the following centiles: 5th, 25th, 50th, 75th, 95th; blue lines indicate male participants, and red lines indicate female participants. Gray overlay reflects kernel density (darker=greater degree of data point overlap). CGC = cingulum (cingulate).

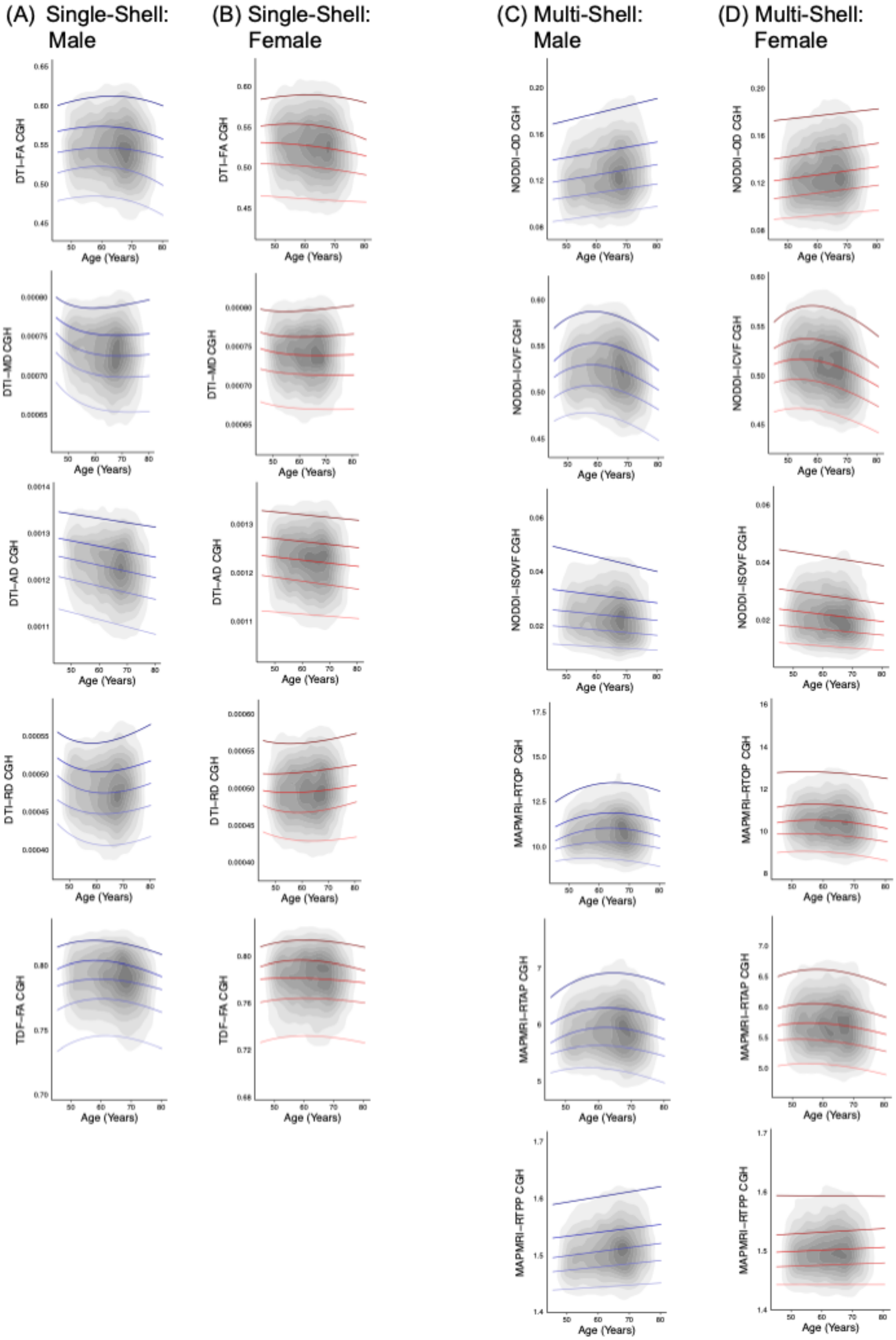


**Figure S3.** Normative centile reference curves calculated for the cingulum (hippocampal) for single-shell dMRI metrics in (A) males and (B) females, and multi-shell dMRI metrics in (C) males and (D) females. Solid colored lines, ordered from lightest to darkest, indicate the following centiles: 5th, 25th, 50th, 75th, 95th; blue lines indicate male participants, and red lines indicate female participants. Gray overlay reflects kernel density (darker=greater degree of data point overlap). CGH = cingulum (hippocampal).

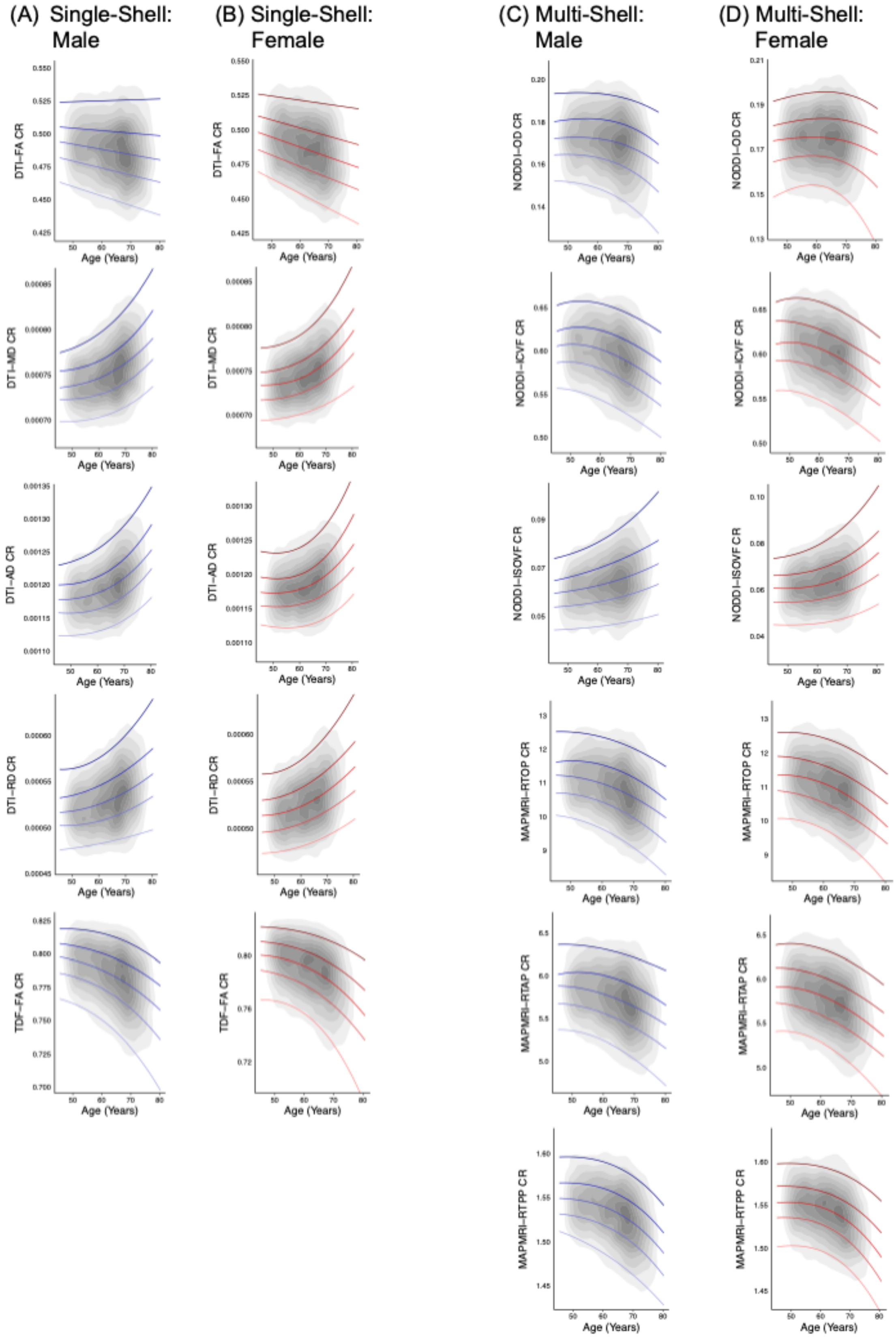


**Figure S4.** Normative centile reference curves calculated for the *corona radiata* for single-shell dMRI metrics in (A) males and (B) females, and multi-shell dMRI metrics in (C) males and (D) females. Solid colored lines, ordered from lightest to darkest, indicate the following centiles: 5$^{th}$, 25$^{th}$, 50$^{th}$, 75$^{th}$, 95$^{th}$; blue lines indicate male participants, and red lines indicate female participants. Gray overlay reflects kernel density (darker=greater degree of data point overlap). CR = *corona radiata*.

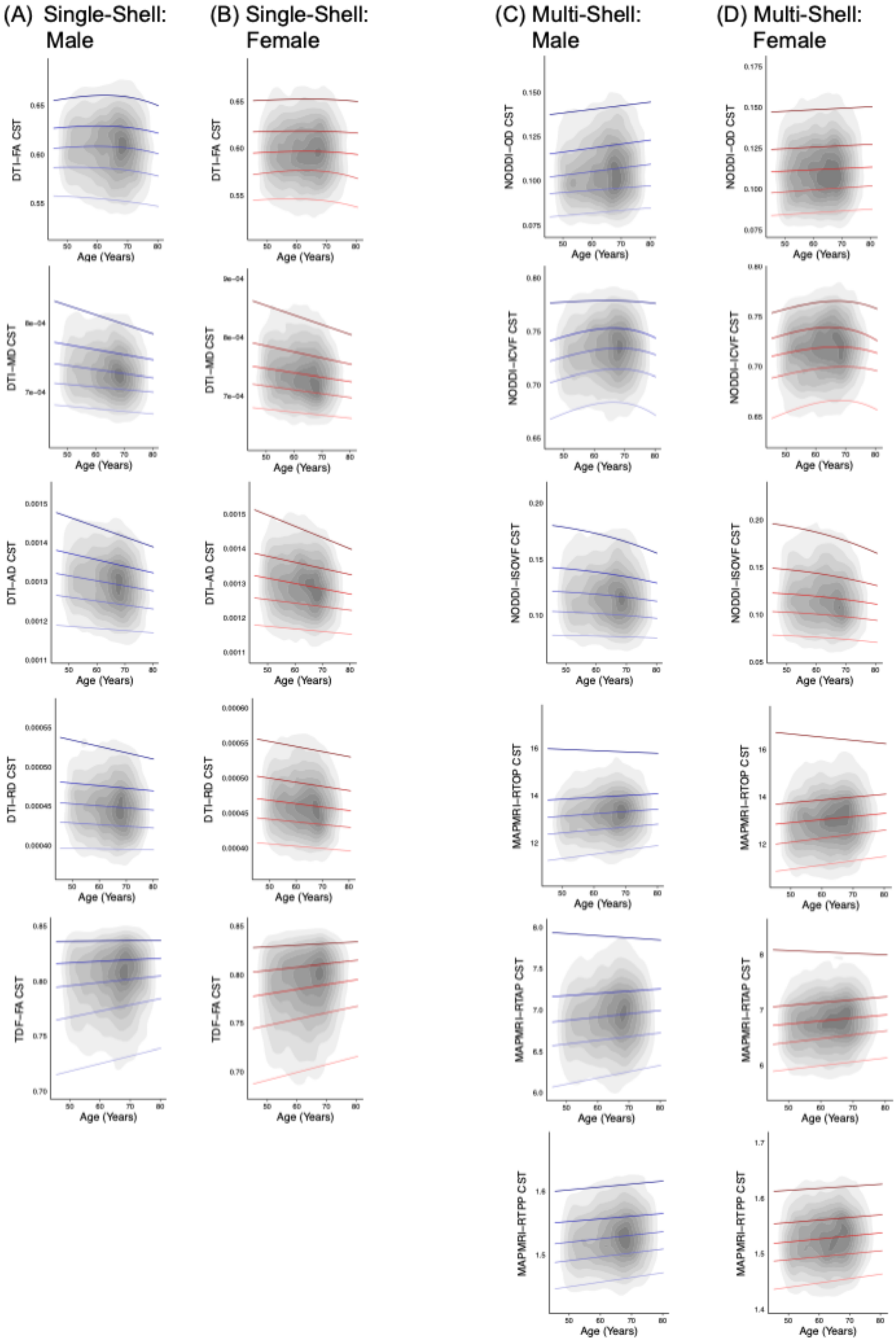


**Figure S5.** Normative centile reference curves calculated for the corticospinal tract for single-shell dMRI metrics in (A) males and (B) females, and multi-shell dMRI metrics in (C) males and (D) females. Solid colored lines, ordered from lightest to darkest, indicate the following centiles: 5th, 25th, 50th, 75th, 95th; blue lines indicate male participants, and red lines indicate female participants. Gray overlay reflects kernel density (darker=greater degree of data point overlap). CST = corticospinal tract.

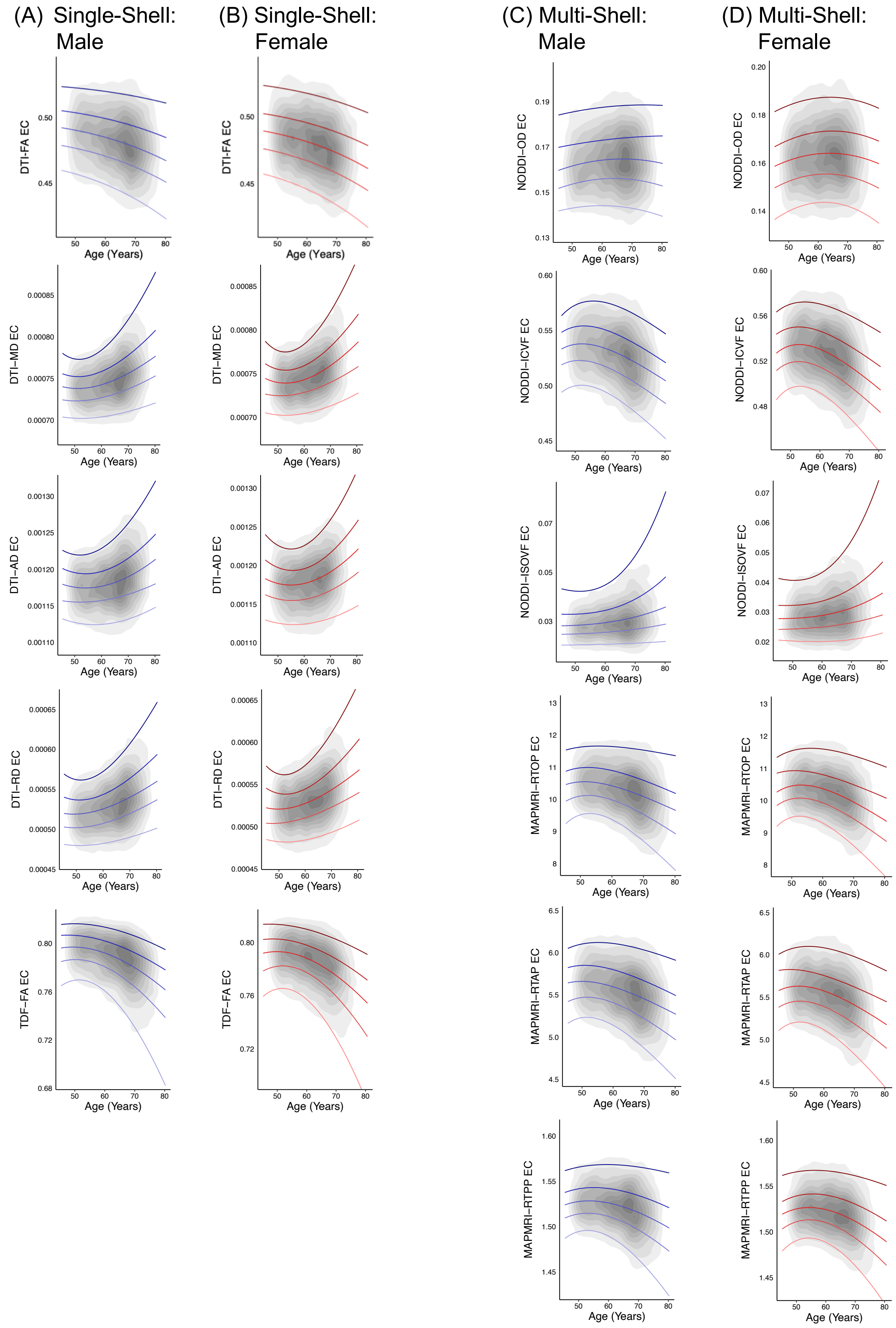


**Figure S6.** Normative centile reference curves calculated for the external capsule for single-shell dMRI metrics in (A) males and (B) females, and multi-shell dMRI metrics in (C) males and (D) females. Solid colored lines, ordered from lightest to darkest, indicate the following centiles: $5^{th}$, $25^{th}$, $50^{th}$, $75^{th}$, $95^{th}$; blue lines indicate male participants, and red lines indicate female participants. Gray overlay reflects kernel density (darker=greater degree of data point overlap). EC = external capsule.

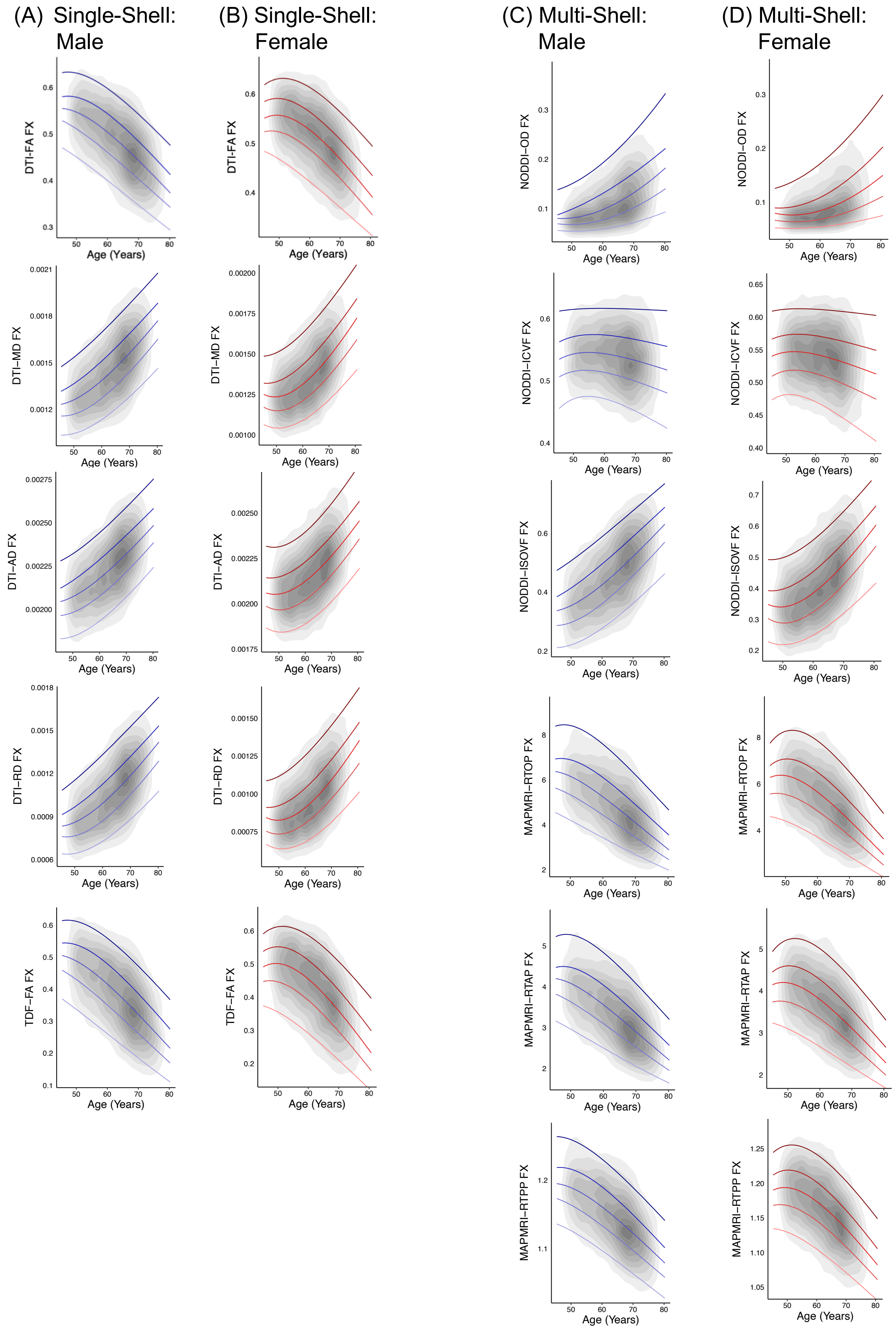


**Figure S7.** Normative centile reference curves calculated for the fornix (body) for single-shell dMRI metrics in (A) males and (B) females, and multi-shell dMRI metrics in (C) males and (D) females. Solid colored lines, ordered from lightest to darkest, indicate the following centiles: 5$^{th}$, 25$^{th}$, 50$^{th}$, 75$^{th}$, 95$^{th}$; blue lines indicate male participants, and red lines indicate female participants. Gray overlay reflects kernel density (darker=greater degree of data point overlap). FX = fornix (body).

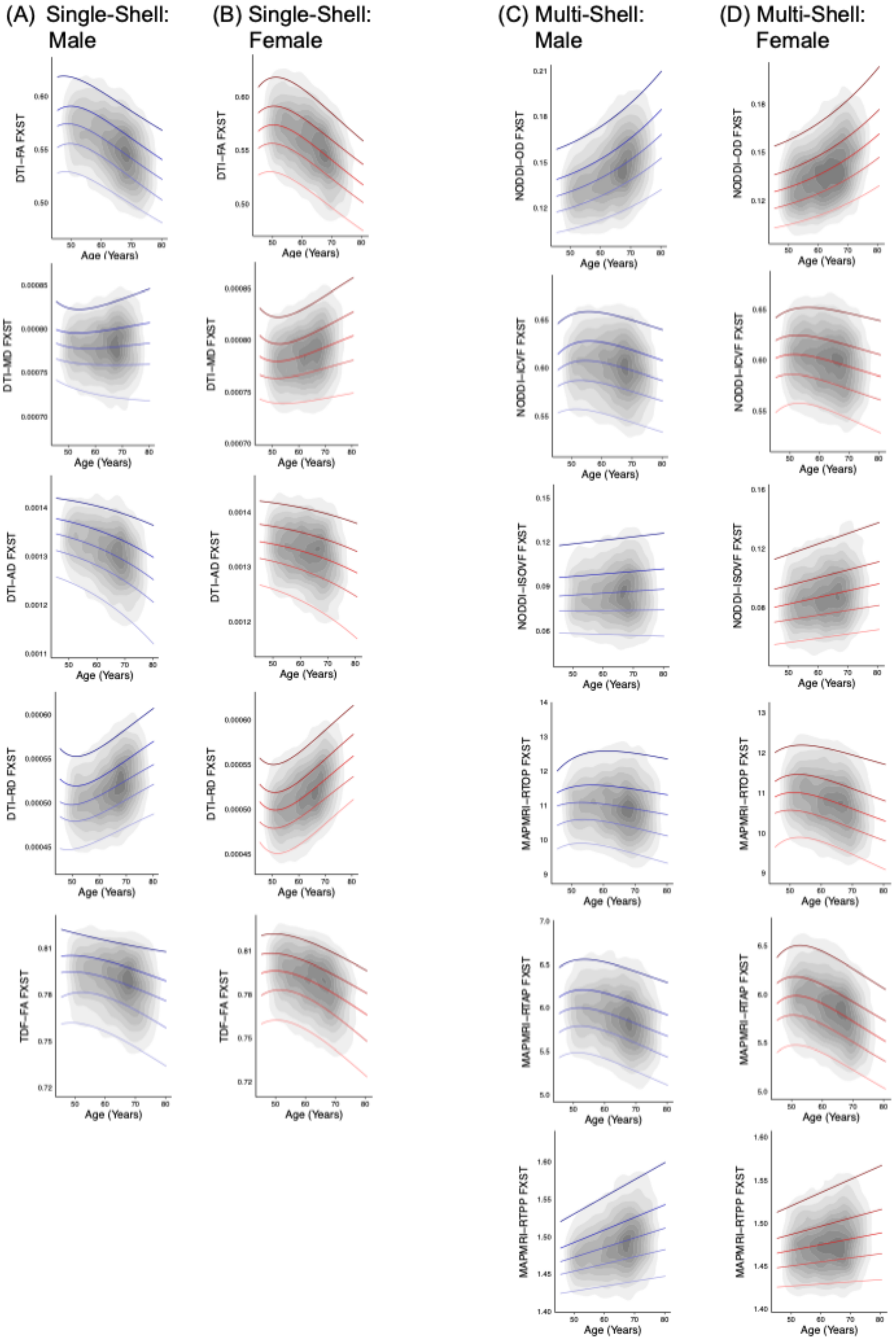


**Figure S8.** Normative centile reference curves calculated for the fornix (*crus*) / *stria terminalis* for single-shell dMRI metrics in (A) males and (B) females, and multi-shell dMRI metrics in (C) males and (D) females. Solid colored lines, ordered from lightest to darkest, indicate the following centiles: 5$^{th}$, 25$^{th}$, 50$^{th}$, 75$^{th}$, 95$^{th}$; blue lines indicate male participants, and red lines indicate female participants. Gray overlay reflects kernel density (darker=greater degree of data point overlap). FXST = fornix (*crus*) / *stria terminalis*.

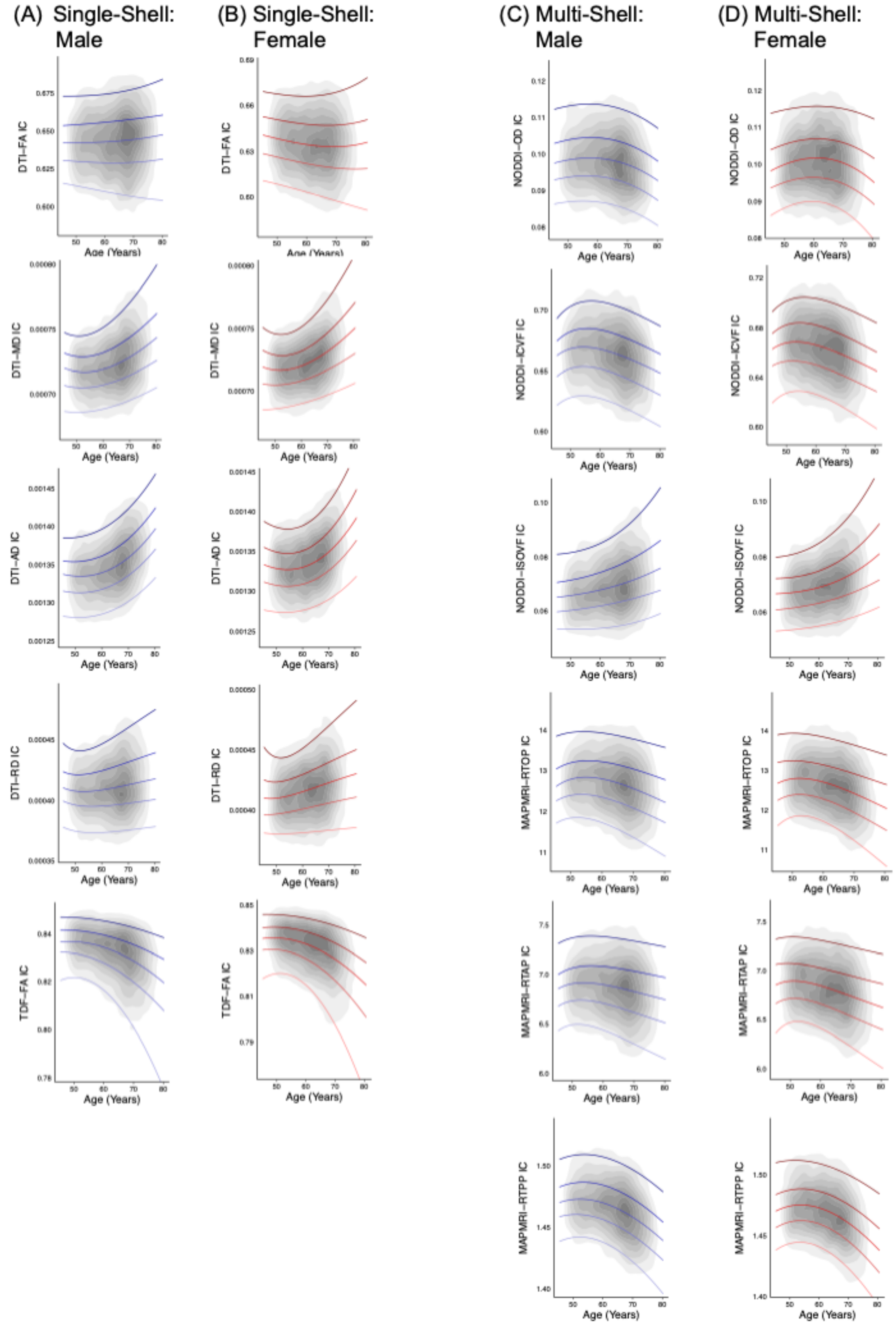


**Figure S9.** Normative centile reference curves calculated for the internal capsule for single-shell dMRI metrics in (A) males and (B) females, and multi-shell dMRI metrics in (C) males and (D) females. Solid colored lines, ordered from lightest to darkest, indicate the following centiles: 5th, 25th, 50th, 75th, 95th; blue lines indicate male participants, and red lines indicate female participants. Gray overlay reflects kernel density (darker=greater degree of data point overlap). IC = internal capsule.

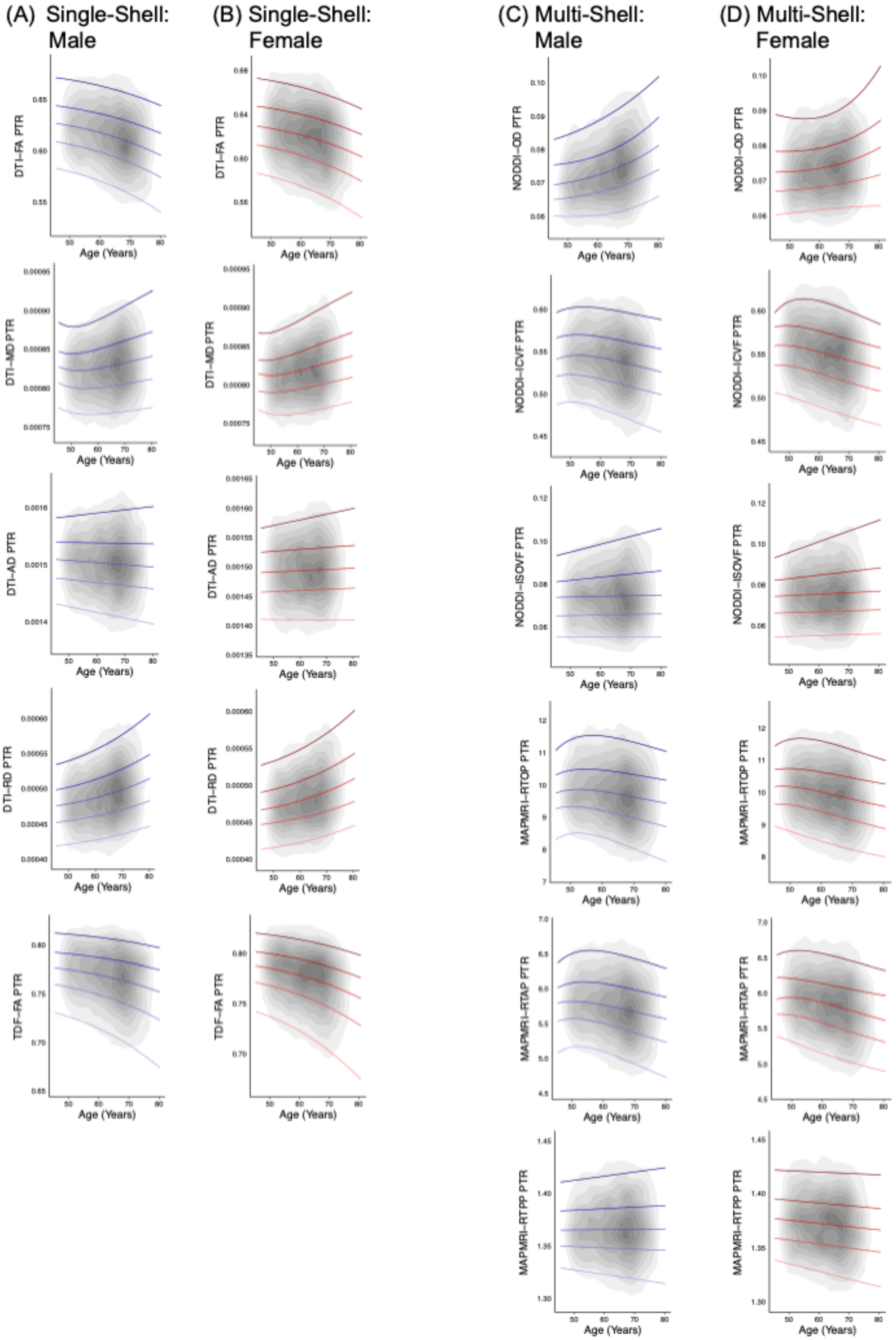


**Figure S10.** Normative centile reference curves calculated for the posterior thalamic radiation for single-shell dMRI metrics in (A) males and (B) females, and multi-shell dMRI metrics in (C) males and (D) females. Solid colored lines, ordered from lightest to darkest, indicate the following centiles: 5$^{th}$, 25$^{th}$, 50$^{th}$, 75$^{th}$, 95$^{th}$; blue lines indicate male participants, and red lines indicate female participants. Gray overlay reflects kernel density (darker=greater degree of data point overlap). PTR = posterior thalamic radiation.

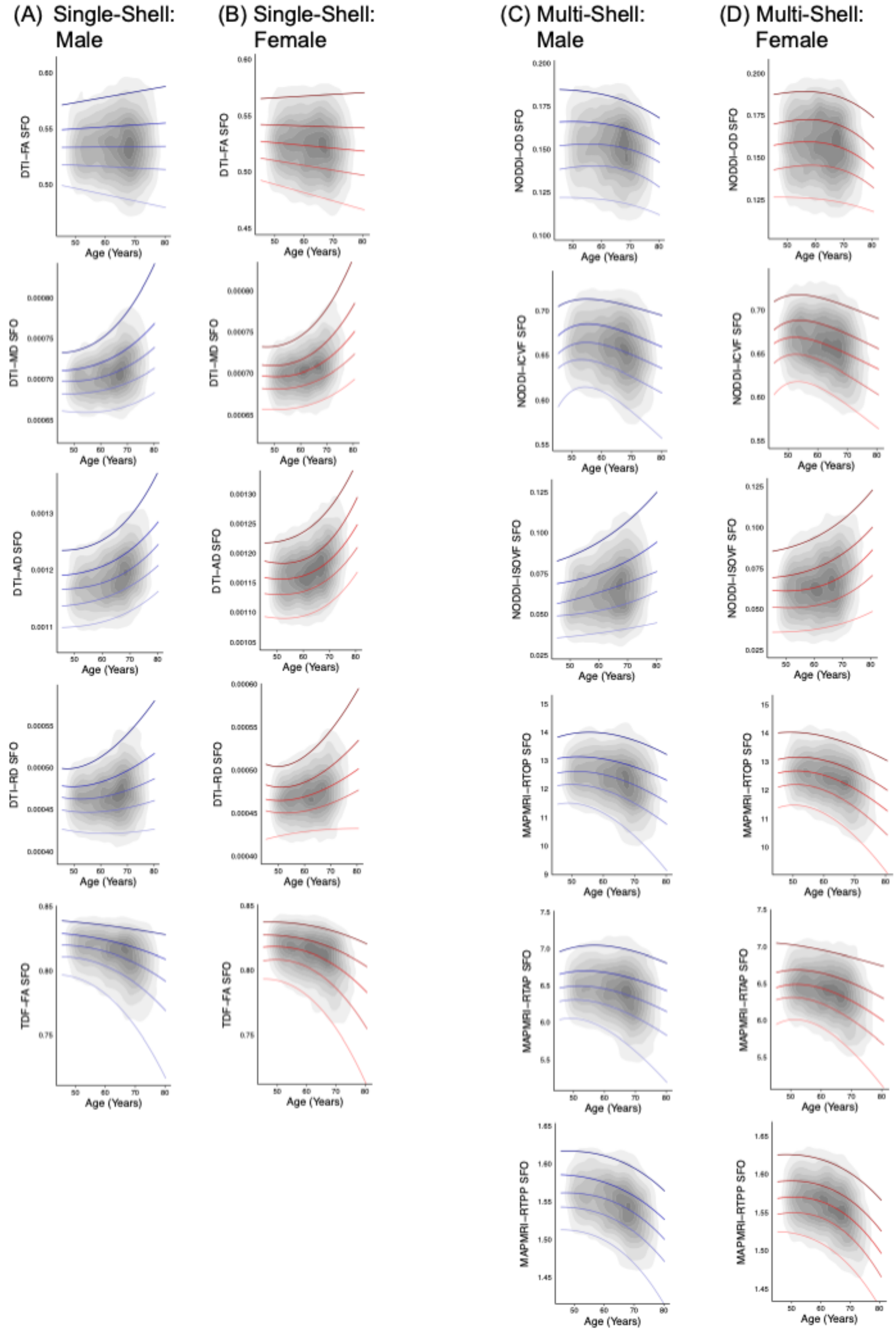


**Figure S11.** Normative centile reference curves calculated for the superior fronto-occpital fasciculus for single-shell dMRI metrics in (A) males and (B) females, and multi-shell dMRI metrics in (C) males and (D) females. Solid colored lines, ordered from lightest to darkest, indicate the following centiles: 5$^{th}$, 25$^{th}$, 50$^{th}$, 75$^{th}$, 95$^{th}$; blue lines indicate male participants, and red lines indicate female participants. Gray overlay reflects kernel density (darker=greater degree of data point overlap). SFO = superior fronto-occpital fasciculus.

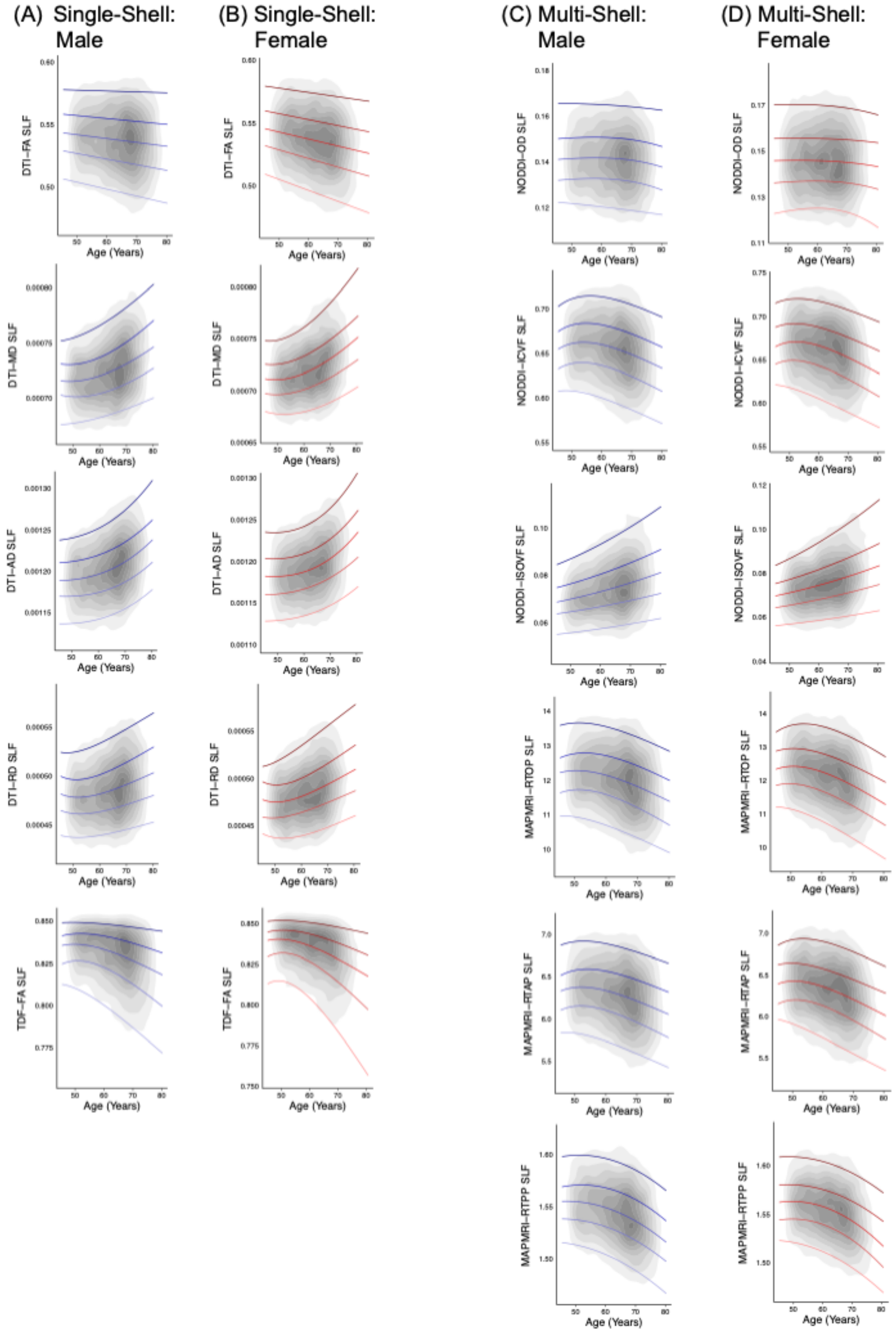


**Figure S12.** Normative centile reference curves calculated for the superior longitudinal fasciculus for single-shell dMRI metrics in (A) males and (B) females, and multi-shell dMRI metrics in (C) males and (D) females. Solid colored lines, ordered from lightest to darkest, indicate the following centiles: 5th, 25th, 50th, 75th, 95th; blue lines indicate male participants, and red lines indicate female participants. Gray overlay reflects kernel density (darker=greater degree of data point overlap). SLF = superior longitudinal fasciculus.

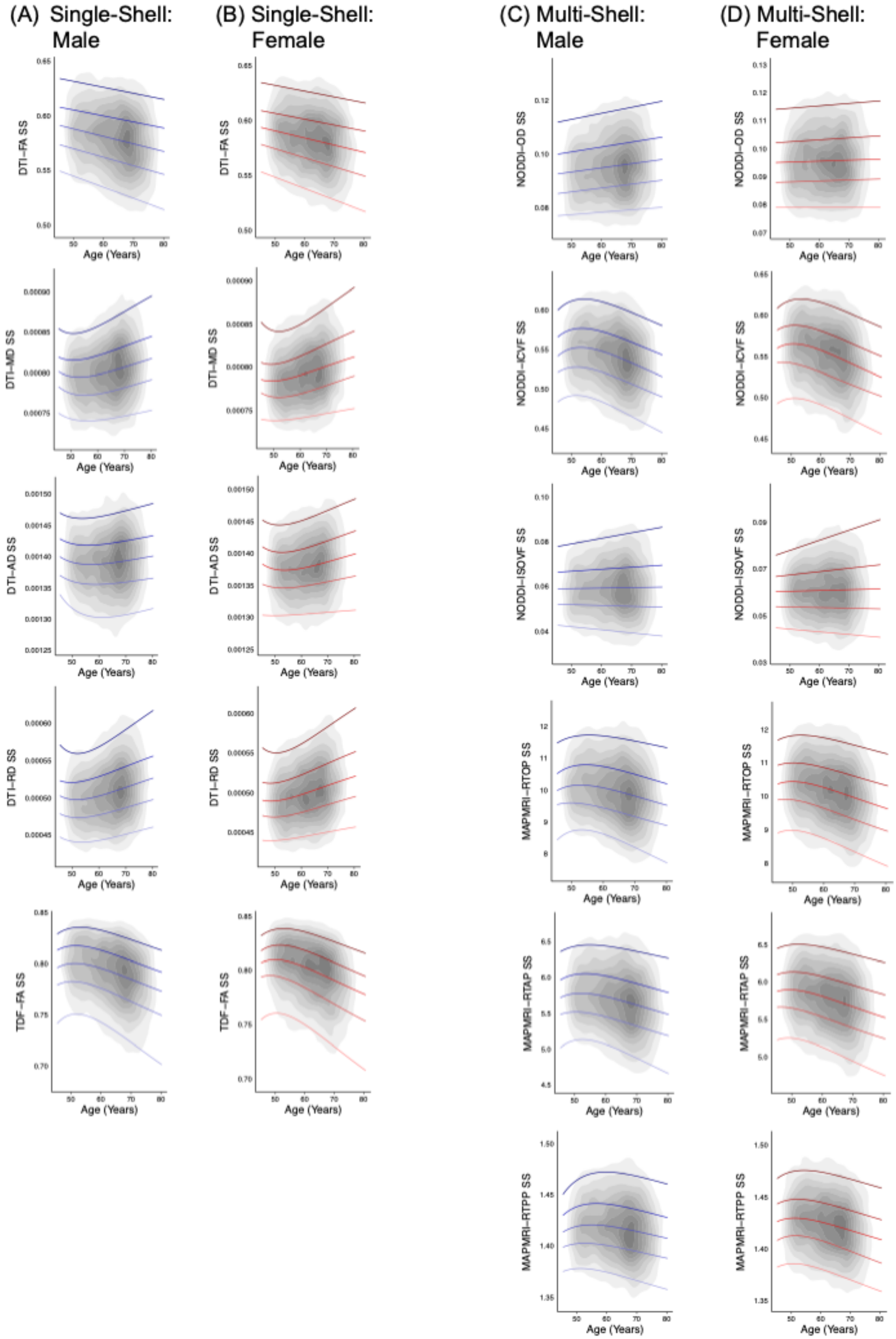


**Figure S13.** Normative centile reference curves calculated for the *sagittal stratum* for single-shell dMRI metrics in (A) males and (B) females, and multi-shell dMRI metrics in (C) males and (D) females. Solid colored lines, ordered from lightest to darkest, indicate the following centiles: 5th, 25th, 50th, 75th, 95th; blue lines indicate male participants, and red lines indicate female participants. Gray overlay reflects kernel density (darker=greater degree of data point overlap). SS = *sagittal stratum*.

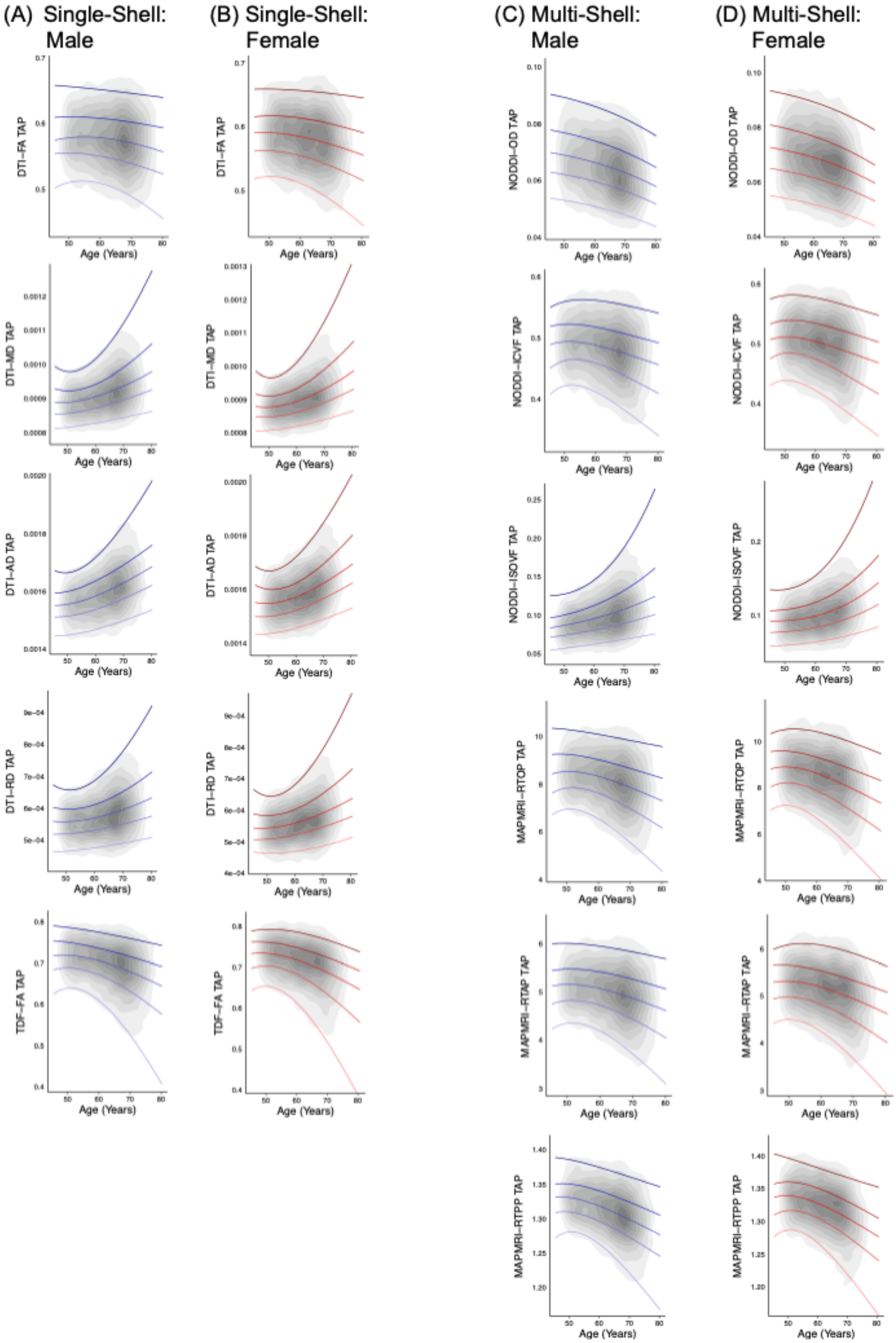


**Figure S14.** Normative centile reference curves calculated for the tapetum for single-shell dMRI metrics in (A) males and (B) females, and multi-shell dMRI metrics in (C) males and (D) females. Solid colored lines, ordered from lightest to darkest, indicate the following centiles: 5th, 25th, 50th, 75th, 95th; blue lines indicate male participants, and red lines indicate female participants. Gray overlay reflects kernel density (darker=greater degree of data point overlap). TAP = tapetum.

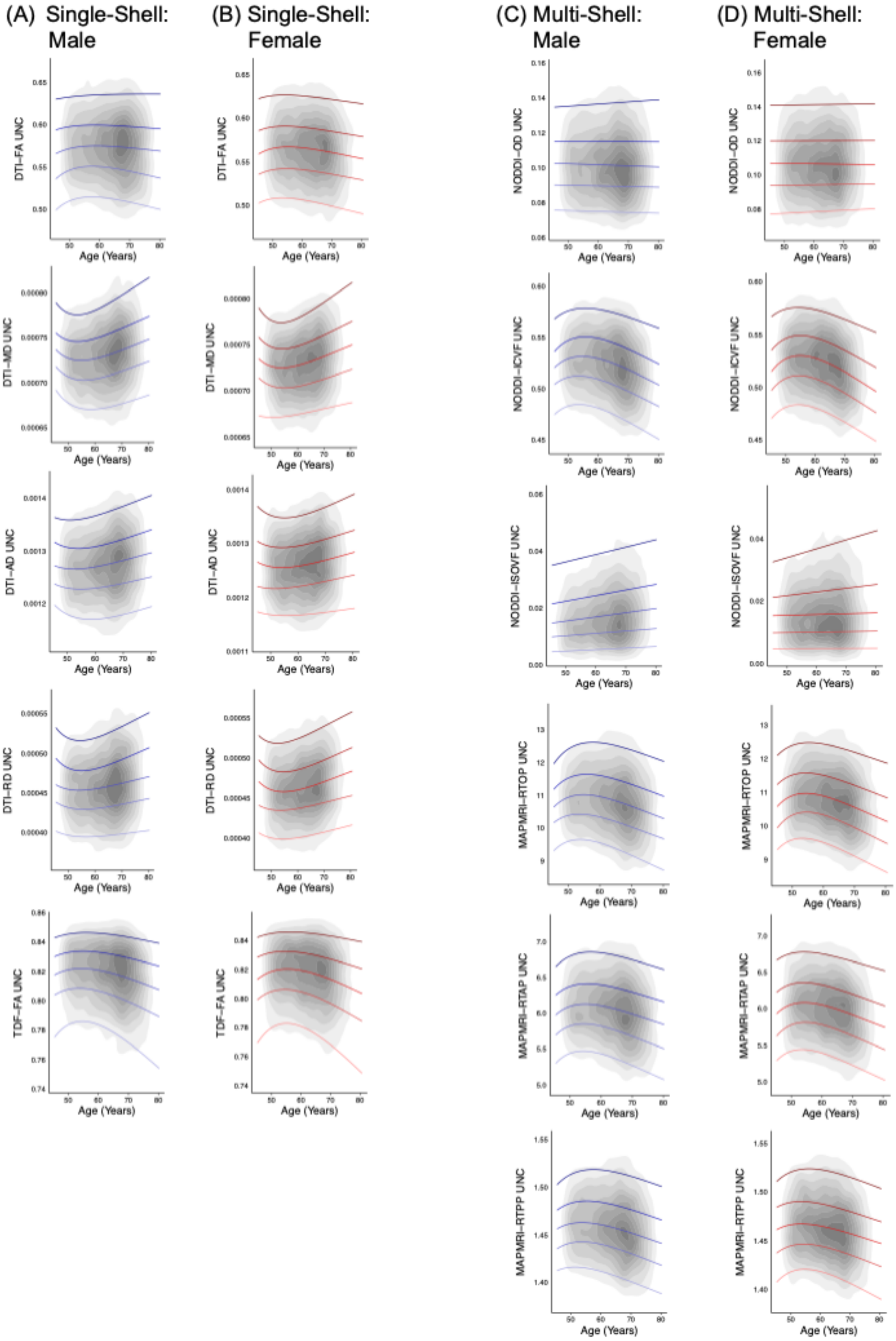


**Figure S15.** Normative centile reference curves calculated for the uncinate fasciculus for single-shell dMRI metrics in (A) males and (B) females, and multi-shell dMRI metrics in (C) males and (D) females. Solid colored lines, ordered from lightest to darkest, indicate the following centiles: 5th, 25th, 50th, 75th, 95th; blue lines indicate male participants, and red lines indicate female participants. Gray overlay reflects kernel density (darker=greater degree of data point overlap). UNC = uncinate fasciculus.

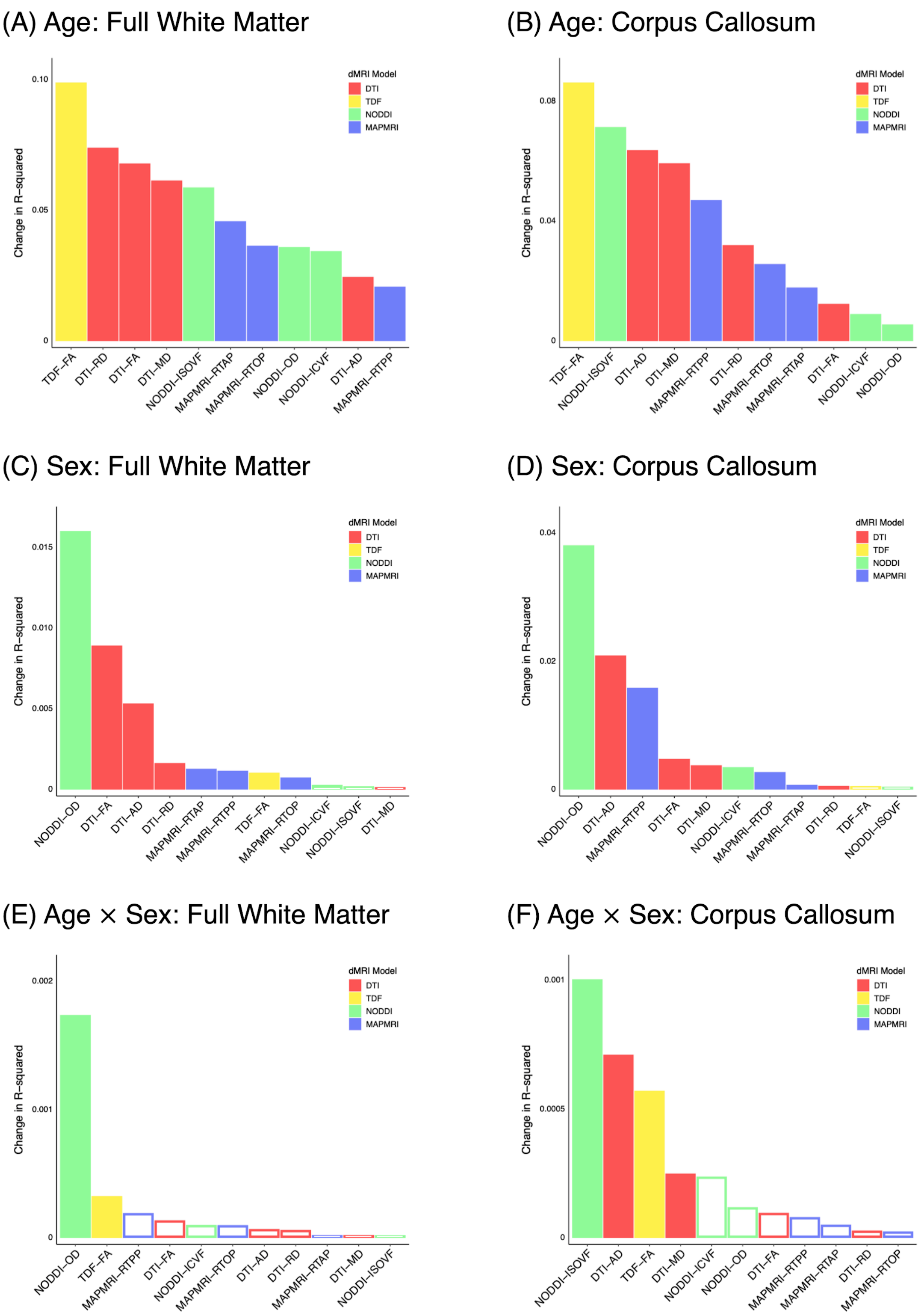


**Figure S16.** Effect of age (A-B), participant sex (C-D), and their interaction (E-F) on full white matter and corpus callosum white matter microstructure. Age was modeled as a discrete variable by splitting participants into two age groups ($\geq$ 60 years or < 60 years). Filled bars indicate a significant association, whereas hollow bars indicate the association did not attain statistical significance.

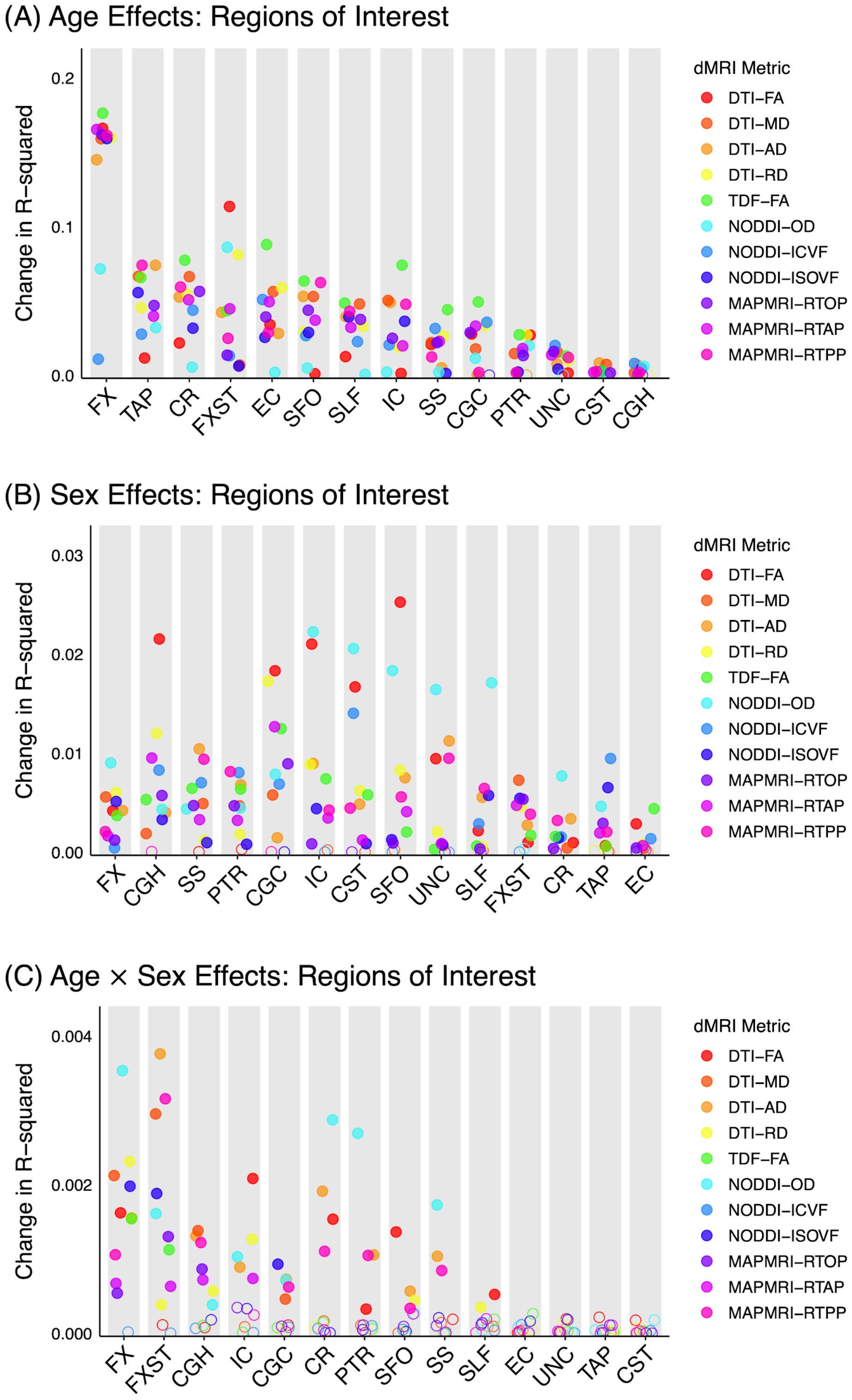


**Figure S17.** Effect of age (A), participant sex (B), and their interaction (C) on white matter microstructure when modeling age by splitting participants into two age groups (≥ 60 years or < 60 years). Filled circles indicate the association was significant after FDR-correction for the number of regions. Regions are ordered by the number of significant metrics, followed by the mean effect size. CGC = cingulum (cingulate), CGH = cingulum (hippocampal), CR = *corona radiata*, CST = corticospinal tract, EC = external capsule, FX = fornix (body), FXST = fornix (*crus*) / *stria terminalis*, IC = internal capsule, PTR = posterior thalamic radiation, SFO = superior fronto-occpital fasciculus, SLF = superior longitudinal fasiculus, SS = *sagittal stratum*, TAP = tapetum, UNC = uncinate fasciculus.
